# GeneTonic: an R/Bioconductor package for streamlining the interpretation of RNA-seq data

**DOI:** 10.1101/2021.05.19.444862

**Authors:** Federico Marini, Annekathrin Ludt, Jan Linke, Konstantin Strauch

## Abstract

**Background:** The interpretation of results from transcriptome profiling experiments via RNA sequencing (RNA-seq) can be a complex task, where the essential information is distributed among different tabular and list formats - normalized expression values, results from differential expression analysis, and results from functional enrichment analyses. A number of tools and databases are widely used for the purpose of identification of relevant functional patterns, yet often their contextualization within the data and results at hand is not straightforward, especially if these analytic components are not combined together efficiently.

**Results:** We developed the `GeneTonic` software package, which serves as a comprehensive toolkit for streamlining the interpretation of functional enrichment analyses, by fully leveraging the information of expression values in a differential expression context. `GeneTonic` is implemented in R and Shiny, leveraging packages that enable HTML-based interactive visualizations for executing drilldown tasks seamlessly, viewing the data at a level of increased detail. `GeneTonic` is integrated with the core classes of existing Bioconductor workflows, and can accept the output of many widely used tools for pathway analysis, making this approach applicable to a wide range of use cases. Users can effectively navigate interlinked components (otherwise available as flat text or spreadsheet tables), bookmark features of interest during the exploration sessions, and obtain at the end a tailored HTML report, thus combining the benefits of both interactivity and reproducibility.

**Conclusion:** `GeneTonic` is distributed as an R package in the Bioconductor project (https://bioconductor.org/packages/GeneTonic/) under the MIT license. Offering both bird’s-eye views of the components of transcriptome data analysis and the detailed inspection of single genes, individual signatures, and their relationships, `GeneTonic` aims at simplifying the process of interpretation of complex and compelling RNA-seq datasets for many researchers with different expertise profiles.

## Background

In modern life and clinical sciences, RNA-sequencing (RNA-seq) is an essential tool for studying gene expression and its regulation (Van den Berge *et al.*, 2019). High-throughput sequencing technologies generate readouts for a large number of molecular entities simultaneously, posing challenges to proper hypothesis generation and data interpretation (Conesa *et al.*, 2016). Among the typical bioinformatic workflows, differential expression (DE) analysis is often employed to identify the genes showing evidence for statistically significant changes, thus being candidate effectors for regulation across the sampled experimental conditions (Love *et al.*, 2015).

Most studies where these techniques are being adopted result in a list containing tens to thousands of gene candidates, with their associated effect size and significance level - often reported as log_2_ fold change (log_2_FC) and adjusted p-values, respectively. Putting these results into biological context by leveraging existing knowledge is essential for facilitating the interpretation of data at a systemic level, and enabling novel discoveries (Chen *et al.*, 2016).

Commonly used knowledge bases for the purpose of functional enrichment analysis include Gene Ontology (GO) (Ashburner *et al.*, 2000; Carbon *et al.*, 2019), KEGG (Kanehisa *et al.*, 2017, 2019), REACTOME (Fabregat *et al.*, 2018), and MSigDB (Liberzon *et al.*, 2011, 2015), where the genes are organized either in simple lists (gene sets, or signatures), or as pathways by accounting for the interactions occurring among the respective members; throughout this manuscript, we will use these terms interchangeably. The analysis at the functional level not only aims to reduce the complexity of high dimensional molecular data (grouping thousands of genes and proteins to just several hundreds of coherent entities), but also increasing the explanatory power of the underlying observed mechanisms (Khatri *et al.*, 2012).

A large variety of computational methods and software have been designed for functional enrichment analysis (Xie *et al.*, 2021), and despite their different implementations, they can still be grouped in three main categories, as identified by Khatri *et al.* (2012): 1. Over-Representation Analysis (ORA), contrasting only the set of DE genes against a background of expressed genes; 2. Functional Class Scoring (FCS), including e.g. Gene Set Enrichment Analysis (GSEA, Subramanian *et al.* (2005)) and its different flavors, incorporating a feature-(gene-)level score/statistic, later aggregated at the pathway level to avoid the choice of a binary threshold; 3. Pathway Topology (PT) based approaches, which utilize the additional information of graph/network structure describing the interactions (Nguyen *et al.*, 2018). The most widely adopted methods in this context are ORA and FCS methods, owing to their ease of applicability, fast runtime, and relevance of resulting gene set rankings, as shown in a recent benchmarking effort (Geistlinger *et al.*, 2020).

Visualization techniques are widely used to make sense of enrichment analysis results, where gene sets might also be highly redundant, thus making the prioritization and interpretation of interesting candidates more challenging (Villaveces *et al.*, 2015; Supek and Škunca, 2017). Numerous tools and applications aim to simplify the interpretation step by adopting a diverse range of methods and visual summaries, and these include BiNGO (Maere *et al.*, 2005), ClueGO (Bindea *et al.*, 2009; Mlecnik *et al.*, 2018), GOrilla (Eden *et al.*, 2009), REVIGO (Supek *et al.*, 2011), GOplot (Walter *et al.*, 2015), AgriGO (Tian *et al.*, 2017), NaviGO (Wei *et al.*, 2017), WebGestalt (Liao *et al.*, 2019), CirGO (Kuznetsova *et al.*, 2019), AEGIS (Zhu *et al.*, 2019), FunSet (Hale *et al.*, 2019), hypeR (Federico and Monti, 2020), KeggExp (Liu *et al.*, 2019), Metascape (Zhou *et al.*, 2019), pathfindR (Ulgen *et al.*, 2019), ShinyGO (Ge *et al.*, 2020), ViSEAGO (Brionne *et al.*, 2019), STRING (Szklarczyk *et al.*, 2019), GSOAP (Tokar *et al.*, 2020), GOMCL (Wang *et al.*, 2020), and netGO (Kim *et al.*, 2020).

Aggregating and summarizing the identified categories is an efficient way to capture and distill the main underlying biological aspects, exploiting visual methods that can efficiently encode the essential information of a table. Among the commonly used visualization methods, many apply different ways of grouping and displaying similar genes or gene sets together, including graph-like representations, clustered heatmaps (either genes by samples, or genes by genesets), or wordclouds. Good visualizations enable discovering underlying trends in the data in an unbiased fashion, and are essential components for the proper communication of results in interdisciplinary projects (Supek and Škunca, 2017; Calura and Martini, 2021).

Datasets and gene set collections increase constantly in their size and complexity, constituting a major barrier for the interpretability of transcriptomic data and their enrichment results, to the point that a potential bottleneck for omics data is the so-called *tertiary* analysis, opposed to mapping and quantification (*primary* analysis) and statistical testing (*secondary* analysis) (Akhmedov *et al.*, 2020). Efficient platforms that enable advanced workflows for a wide range of users can play a big role in providing the required level of interactivity, while guaranteeing the adherence to gold standard methods and to best practices for reproducible analyses (Sandve *et al.*, 2013; Marini and Binder, 2016; Brito *et al.*, 2020).

The different atomic elements for a typical RNA-seq analysis (expression table, results from differential expression, functional enrichment results) can stem from different pipeline outputs, yet they need to be combined together, e.g. in a report created following the rules of literate programming (Knuth, 1984). By providing accessible summaries with proper data visualization and interpretation methods, in formats that facilitate dynamic shareable outputs, such frameworks can greatly reduce the time to generate novel hypotheses and insight. Often, this task is not straightforward to carry out, as different software solutions or environments might be chosen, resulting in different file formats, thus increasing the difficulty for practitioners to explore all relevant aspects of the data at hand, even if common sets of gene and pathway identifiers are adopted.

A number of solutions have been developed in diverse languages (mostly R, Python, Java) to address the challenges listed above, but no software package provides a comprehensive framework for assisting the proper interpretation of RNA-seq data; interested readers can find a comparative overview of the features of the above mentioned tools in Suppl. Table S1.

Here we present `GeneTonic`, an R/Bioconductor package aiming to streamline the identification of relevant functional patterns, as well as their contextualization in the data and results at hand, by combining in a seamless way all the pieces of information relevant for a transcriptomic analysis. The `GeneTonic` package is composed by a Shiny web application, with a variety of standalone functions to perform the analysis both interactively as well as in a programmatic way. `GeneTonic` requires as input the results generated by each analytic step (quantification, DE testing, functional enrichment), which are usually shared as separate tables or spreadsheets by bioinformaticians and core facility service providers, in formats that are suitable to standardization.

`GeneTonic` makes it easy to generate visualizations, starting from bird’s eye perspective summaries (gene-geneset graphs, enrichment maps, also linked to interactive tables in the web application), as well as getting in depth dedicated summaries for each geneset of interest. User actions enable further insight and deliver additional information (e.g. gene info boxes, geneset summaries, and signature heatmaps), with drilldown tasks activated by simple mouse clicks. While simple operations within the call to the `GeneTonic()` main function makes the result set more interpretable, our package also supports built-in RMarkdown reporting as a foundation for computational reproducibility, to conclude an interactive exploration session (Marini and Binder, 2019; Marini *et al.*, 2020). We carefully designed the user interface, enabling the required tasks in a straightforward way, as a result of an open and continuous dialogue with researchers adopting this tool in its early development. Users can learn-by-doing the functionality of `GeneTonic` via guided tours, creating a common ground for experimentalists and analysts to explore transcriptomic data at the desired depth and efficiently generate novel insights (Poplawski *et al.*, 2016).

`GeneTonic` connects together a number of R/Bioconductor packages, implementing the current best practices in RNA-seq data analysis, and facilitates the communication between experts of different disciplines. Harmonizing the output of the many analysis steps, possibly performed also with a variety of tools, `GeneTonic` is a powerful tool for digesting and enjoying any RNA-seq dataset: the interactivity is a compelling means to empower end users for the exploration of many features of interest, and by providing a report with full code snippets, we support analyses that are reproducible and easily extendable. The `GeneTonic` package is available at https://bioconductor.org/packages/GeneTonic/, and a public instance is available for demonstration purposes at http://shiny.imbei.uni-mainz.de:3838/GeneTonic.

## Implementation

### General design of GeneTonic

The `GeneTonic` package is written in the R programming language, leveraging many existing packages currently available in the Bioconductor project, which constitute the foundation for a broad spectrum of analytic workflows in computational biology and bioinformatics (Huber *et al.*, 2015; Amezquita *et al.*, 2019), and the Shiny framework for interactivity (Chang *et al.*, 2020). The typical use case for `GeneTonic` expects researchers to run the web application locally, providing the atomic components of a typical RNA-seq analysis workflow (Fig. 1, top section).

**Figure 1:**
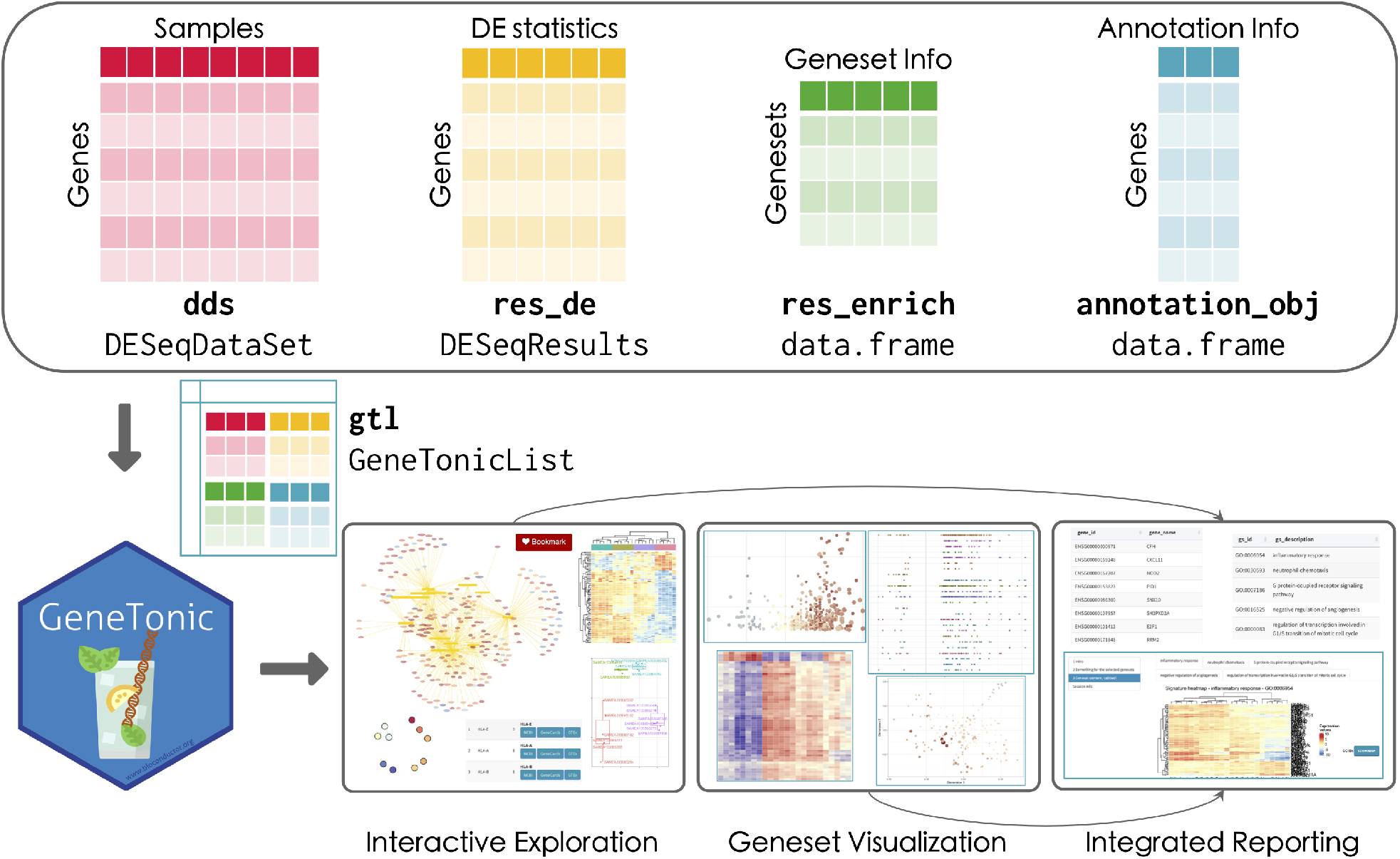
Overview of the GeneTonic workflow. Four elements are required to optimally use the functionality of `GeneTonic` (top section): dds, a `DESeqDataSet`, with the information related to the expression values; `res_de`, a `DESeqResults` object, storing the results of the differential expression analysis; res_enrich, a data frame with the result from the enrichment analysis; annotation_obj, a data frame containing different sets of matched identifiers for the features of interest. These can be combined into a `GeneTonicList` object (middle section), which can be directly fed to `GeneTonic` and its functions, enabling (bottom section) interactive exploration and a variety of visualizations on the geneset enrichment results, both exploited to generate an integrated HTML report of the provided data.

`GeneTonic` is designed to be used after the main steps of DE and functional enrichment analyses have already been completed. While this might seem a limiting factor, we wanted to acknowledge that a plethora of validated methods for performing functional analyses at the pathway level exist, and similarly, well established statistical methods for DE are available (Geistlinger *et al.*, 2020; Van den Berge *et al.*, 2019). Our focus was rather on providing a standardized interface (via so-called shaker functions) to automatically handle the outputs of the different tools which most users might be familiar with, so that `GeneTonic` retains a wide applicability with respect to the upstream analysis workflows - this is illustrated in the use cases of Suppl. Files 1 and 2.

The required input components are stored as in the `DESeq2` workflow (Love *et al.*, 2014), using classes descending from the versatile `SummarizedExperiment` container, *de facto* the adopted standard for interoperability in the Bioconductor ecosystem (Huber *et al.*, 2015). Tabular information can be provided as simple data frame objects, either imported from textual output of the different tools, or converted internally by the `shaker` functions of GeneTonic. We encourage users to adopt stable feature identifiers, such as ENSEMBL or Gencode (Yates *et al.*, 2019; Frankish *et al.*, 2019), and enable the automated conversion to HGNC gene symbols via annotation tables.

Most of the implemented functionality can be accessed by a single call to the main `GeneTonic()` function, with the different visualizations and summaries directly available from the dedicated sections of the web application. These functions are also exported for usage in scripted analyses such as RMarkdown HTML reports, making it easy to automate tasks while still creating interactive widgets that can be explored in depth offline (Fig. 1, bottom section).

The user interface (shown in Fig. 2) is structured with the layout provided by the `bs4Dash` package (Granjon, 2019), which implements Bootstrap 4 over the infrastructure of `shinydashboard` (Chang and Borges Ribeiro, 2018). The main features include a sidebar menu (Fig. 2A) to navigate the different sections of the app, a header and a collapsible control panel to provide widgets which define the general behavior of the main panel, displayed in the dashboard body (Fig. 2C).

**Figure 2:**
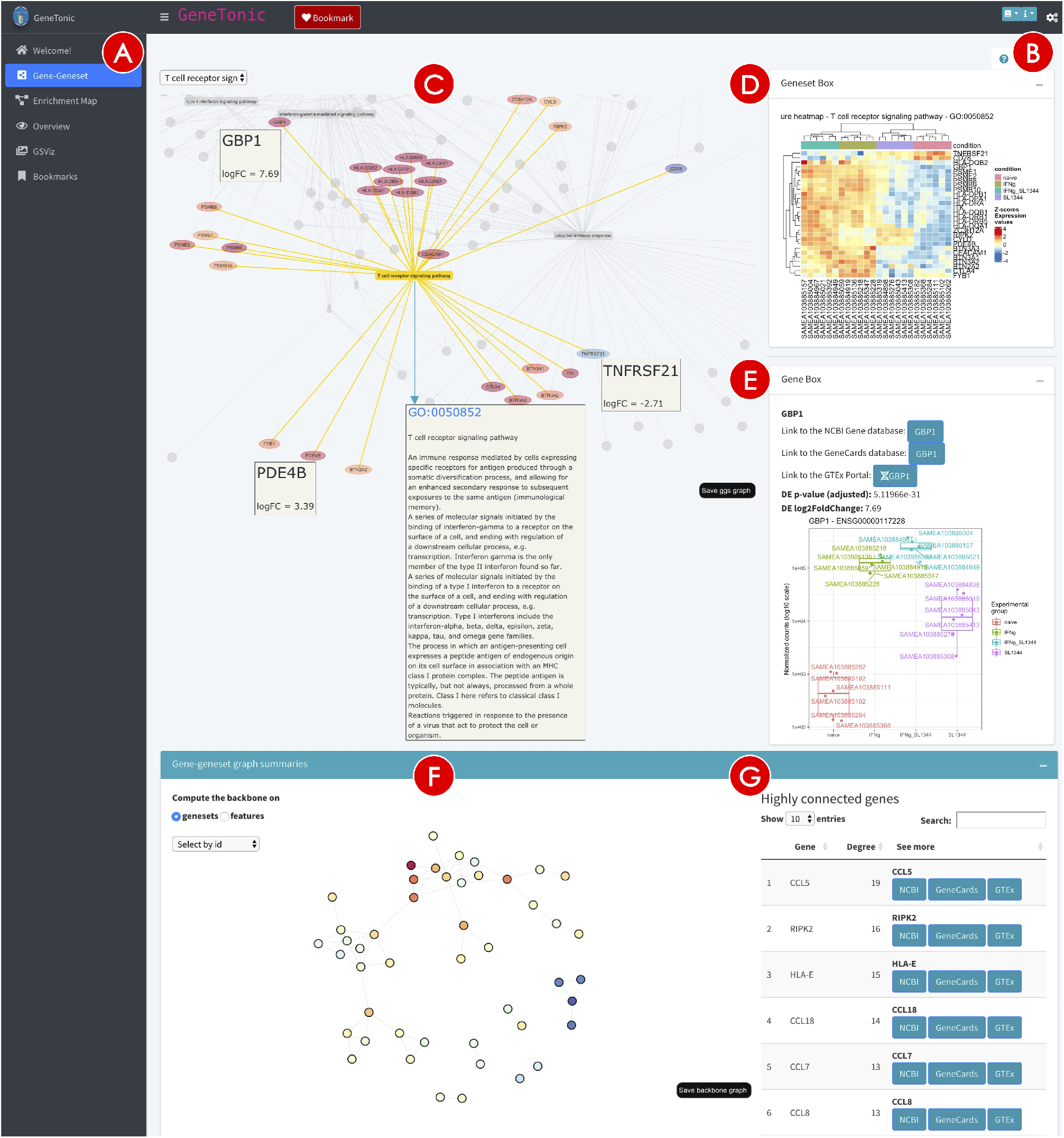
Screenshot of the *Gene-Geneset* panel in the GeneTonic application. The main navigation of the app is performed via the sidebar menu (**A**), while the options common to the different panels are available in the control bar (toggled with the cogs icon, close to the **B** circle). The main area of the *Gene-Geneset* panel (**C**) contains an interactive bipartite graph connecting genesets to their respective components, color coded according to the type and the expression changes. Upon clicking on any node, a Geneset Box (**D**) or a Gene Box (**E**) can be automatically generated for facilitating further exploration. The backbone of the graph above (**F**) represents a compact summary of the bipartite projections, and a table displays the most highly connected nodes (**G**), as potential hubs with regulatory roles in multiple genesets.

To instruct users on how to efficiently leverage the exploration components, we enhance the content provided in the use case vignette with guided tours of the interface (Fig. 2B), implemented via the `rintrojs` package (Ganz, 2016). This learning-by-doing paradigm invites the user to perform the actions that reflect typical usages in each module, and can be seen as a dynamic extension of the static documentation format.

Collapsible and tab-based elements allowed us to build a rich user interface, yet without adding too much visual clutter, which would hamper the usability of the analysis sessions - and by that reduce the ability to extract relevant insight.

All the required elements for running `GeneTonic` are provided at the beginning of the execution, meaning that the navigation throughout the different modules can take place with the usual iteration cycles that build up a full in-depth exploration. As this process can become time-consuming, we implemented a dedicated bookmarking system to temporarily store the genes and gene sets of interest, either by clicking on the dedicated button or with a keystroke (defaulting to the left control key). A summary for these selected features is automatically rendered in the *Bookmarks* section, where the user can generate a full report on the provided input parameters, focusing on the aspects picked up during the live session (Fig. 1, bottom). It is then easy to reconstruct and reproduce the analytic rationale, and share the rendered outputs with cooperation partners, or simply store them for the purpose of careful documentation.

### Typical usage workflow

The typical session with `GeneTonic` can start once the required inputs are provided to the main function (as illustrated in Fig. 1).

In order to use `GeneTonic`, the following inputs are required: 1. `dds`, a `DESeqDataSet`, the main component in the `DESeq2` framework, storing the information related to the expression matrix; 2. `res_de`, encoded as `DESeqResults` for containing the results of the differential expression analysis; 3. `res_enrich`, i.e. the result from the enrichment analysis, likely converted through one of the shaker functions for preprocessing (or manually, if feeding this from a tool currently not supported), structured as a data frame with a minimal set of required variables (pathway identifier, description, significance level, and affected genes); 4. `annotation_obj`, the gene annotation data frame, i.e. a table with at least two columns, `gene_id` for a set of unambiguous identifiers (e.g. ENSEMBL ids), and `gene_name`, containing a human-readable set of names, e.g. HGNC-based gene symbols.

Conveniently, a single named list containing these inputs (Fig. 1, middle section) can be provided as an alternative format, with many functions of `GeneTonic` accepting a `gtl` parameter (standing for “`GeneTonicList`”). This simplifies the creation of context-dependent serialized objects that can be easily shared by data analysts to experimental collaborators.

In its current version, `GeneTonic` can directly handle the output of different tools, selected for being among the most commonly used in pathway analysis, including `topGO` (Alexa *et al.*, 2006), `clusterProfiler` (Yu *et al.*, 2012), DAVID (Huang *et al.*, 2009), Enrichr (Kuleshov *et al.*, 2016), g:Profiler (Reimand *et al.*, 2019; Raudvere *et al.*, 2019), and `fgsea` (Korotkevich *et al.*, 2021) - these are showcased in the code included in Suppl. Files 1 and 2. We plan to extend the compatibility of `GeneTonic` with the output of newly developed tools, or alternatively welcome contributions on the project homepage on GitHub (https://github.com/federicomarini/GeneTonic).

All the components of the `GeneTonic()` application can be seamlessly used by leveraging sets of shared gene identifiers across the different input objects. This makes it possible to compute aggregate scores for each gene set, e.g. averaging the log_2_ fold change of all the affected members of the gene set, or computing a Z-score based on the standardized sum of the number of genes regulated in either direction. As gene sets cannot take into account the topological information of a pathway, this is a valid surrogate means to summarize the effect between conditions at the functional level, and can be visually encoded in the outputs of the `GeneTonic` dedicated routines.

The process of data exploration and interpretation is iterative by its own nature. `GeneTonic` supports this by employing a variety of visual summaries (gene-geneset bipartite graphs, enrichment maps, geneset volcano plots, and more in the dedicated app sections - Fig. 2 and Suppl. Fig. 1). We also offer methods to efficiently extract the most meaningful affected biological themes, e.g. by grouping similar categories and selecting a representative pathway for each subset, in order to simplify the redundancy often found in functional enrichment results. Whenever possible, we provide additional information boxes for genes and genesets (Fig. 2D and E) to facilitate drilldown tasks and better understand the whole data components of the project. A number of automatically generated action buttons link directly to external databases, such as AmiGO (Carbon *et al.*, 2019), NCBI (Agarwala *et al.*, 2017), GeneCards (Stelzer *et al.*, 2016), GTEx (Gamazon *et al.*, 2018), enabling more in-depth analysis of particular genesets or genes, without the need to type all the entries of interest.

While the main way of using the functionality of `GeneTonic` is probably via its web application, we designed all the underlying functions to be able to handle standard objects and classes adopted by the current Bioconductor workflows, and therefore their output can be also incorporated in information-rich HTML reports and existing scripted analyses without additional effort. Indeed, the report itself created via the `happy_hour()` function is an exemplary RMarkdown document, which users can edit and extend as they see fit. Literate programming approaches were initially conceived by the seminal work of Knuth (Knuth, 1984) and have been currently refined in the knitr framework (Xie, 2013) and in the Jupyter notebook system (Rule *et al.*, 2019). These techniques constitute a powerful toolkit to ensure the reproducibility of computational analyses (Sandve *et al.*, 2013; Stodden and Miguez, 2014; Brito *et al.*, 2020). The creation of such an HTML document is also intended as the recommended concluding step of a typical usage session for `GeneTonic`.

In case additional or bespoke visual representations of the input objects (e.g. MA plot for the DE results, customized heatmaps, …) should be created, the `iSEE` Bioconductor package (Rue-Albrecht *et al.*, 2018) can be used for this purpose. We provide a specific export function, combining the provided inputs into a `SummarizedExperiment` object that can be readily further explored in an instance of `iSEE`, by properly accessing the `assays`, `colData`, and `rowData` slots.

## Results

In this section, we will illustrate the functionality of `GeneTonic`, showcasing the results for the analysis of a human RNA-seq dataset of `macrophage` immune stimulation, published in Alasoo *et al.* (2018). The data is made available via the macrophage Bioconductor package, which contains the files output from the Salmon quantification (version 0.12.0 - Patro *et al.* (2017)), against the Gencode v29 human reference. Expression values, summarized at the gene level, are available from 6 individual donors, in 4 different conditions. We will focus on the comparison between Interferon gamma treated samples vs naive samples.

A comprehensive report on the processing for this dataset and its usage with `GeneTonic` is included in Suppl. File 1. Suppl. File 2 showcases the usage of `GeneTonic` on the findings of the work of Mohebiany *et al.* (2020) (A20-deficiency in mouse microglia cells), describing also an alternate entry point for running `GeneTonic` with objects from the `edgeR` workflow for differential expression (Lun *et al.*, 2016).

### Augmenting functional enrichment results with expression data

The majority of functions in `GeneTonic` requires only a minimal set of information on the pathways enrichment results, i.e. a gene set identifier, its description, and a measure of significance for the enrichment, often specified as a p-value. Nevertheless, it is often beneficial to combine additional knowledge, if provided by the method used to perform the enrichment test; this might include the number of genes annotated to each pathway and the related subset detected as differentially expressed, but more importantly it can incorporate the full set of expression values in the original dataset.

One way to do so is via the `get_aggrscores()` function, which computes the overall pathway Z-score and an aggregated score (such as the mean or the median), which summarize at the gene set level the effect (log_2_FC) of the differentially expressed genes annotated as its members. In detail, the gene set Z-score attempts to determine the “direction” of change, and is computed as 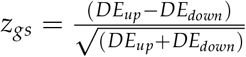, where *DE_up_* and *DE_down_* are the number of up-regulated and down-regulated DE genes annotated to the geneset *gs*, respectively (Fig. 3A and B).

**Figure 3:**
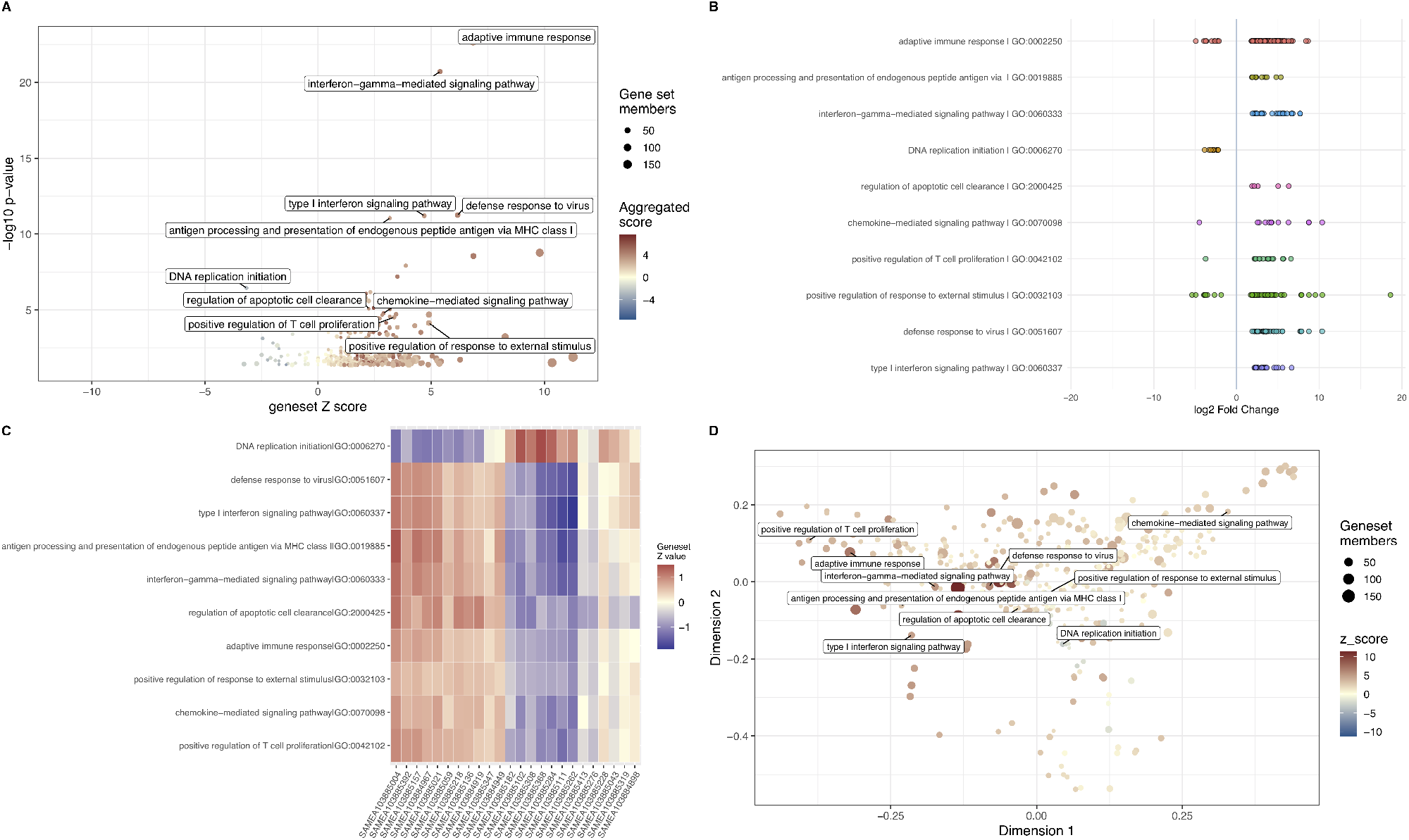
Graphical summaries of enrichment results. **A** A geneset volcano plot (`gs_volcano()`), with the −log_10_p-value depicted against the geneset Z-score, with a subset of representative affected functions, generated from the output of the `gs_fuzzyclustering()` function. The size of the points maps the information of the geneset size. **B** An enhanced visual summary for the enrichment results, displaying the contributions of the single genes to each gene set (with the directionality as log_2_FC), created with `enhance_table()`. **C** A heatmap for the matrix of sample-wise geneset Z-scores for the same subset used in the other subpanels, generated with `gs_scoresheat()` on the output of `gs_scores()`. **D** A multidimensional scaling plot for genesets, colored by their Z-score, representing the set similarity. This can be generated with the `gs_mds()` function.

Alternatively, a sample-level gene set score can be computed, in an approach similar to the implementation of `GSVA` (Hänzelmann *et al.*, 2013). First, a variance stabilizing transformation is applied to the expression matrix, returning values that show a higher degree of homoscedasticity, thus more amenable to downstream processing and visualization. For each gene, the values are Z-standardized by subtracting the row-wise mean and dividing by the row-wise standard deviation.

Finally, for each pathway, we take the subset of the Z values corresponding to its members, and its average is computed and returned as pathway score. We define *Z_ij_* as the Z-score for gene *i* in sample *j* as 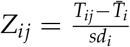, whereas *T_ij_* is the correspondent entry in the transformed expression values matrix, 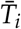 and *sd_i_* are the mean and standard deviation for the gene *i*, respectively. The entry *GS_kj_* for pathway *k* in sample *j* is thus defined as 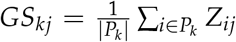, where *P_k_* is the set of DE genes for pathway *k* (Fig. 3C).

This extra information about the status of activation/repression for each pathway can be efficiently encoded as aesthetic elements in plots (e.g. the color of a node in a graph, with the geneset Z-score), or directly displayed as a heatmap of the pathway score matrix to compare the activity among the samples.

### Exploring the interplay of pathways and genes, interactively

The relationships among pathways and their member genes, or just between different pathways, can quickly become hard to manage when using simple textual or tabular formats. This can be due to the growing size of existing annotations, whereas the increase in detail can also lead to an increase in redundancy, thus making the task of extracting the key biological messages harder.

A number of visualization techniques have been adopted in the last years to simplify this basic yet essential operation (Supek and Škunca, 2017; Calura and Martini, 2021), and a common way to represent this complex interplay is by using graphs. Unipartite graphs are an efficient way to depict the degree of similarity among genesets, where genesets themselves are the nodes, and edges encode for information such as the degree of similarity/overlap between the two nodes (Merico *et al.*, 2010) - see Suppl. Fig. 1. Bipartite graphs (as in Fig. 2) can be naturally adopted to include both genes and genesets as the main node types, with unweighted edges representing in this case the binary membership status for one gene with respect to one geneset (Pomaznoy *et al.*, 2018).

`GeneTonic` builds upon these foundations and implements the possibility to interact with the nodes upon hovering with the mouse (or clicking on them). The graph objects are generated dynamically, including the desired number of genesets; by default, the top most significant hits in the enrichment results are selected. Interactivity is provided by the `visNetwork` package (Almende B.V. *et al.*, 2019), that wraps the vis.js library bindings, building on the `htmlwidgets` framework. Depending on the type of node selected in the main user interface, an information box is populated (Fig. 2D and E).

If selecting a pathway (displayed as a yellow box), the info box will contain details on the geneset (if detected as a Gene Ontology term), and a signature heatmap is displayed, with the variance stabilizing transformed expression data encoded as color to give a fine-grained view of the behavior for all its set members; this is particularly useful to connect the existing biological background with lists of features where no information on the topology is provided, enabling to detect subgroups of correlated expression patterns. Another useful representation can be obtained by coupling a volcano plot (for representing differential expression) with the annotated labels of the members of a geneset; this is implemented in the `signature_volcano()` function, and displayed in the same info box (Suppl. Fig. 1).

Genes are displayed in the graph as ellipses, colored using a divergent palette to encode for the effect size as log_2_ fold change; when a gene is selected, a plot for the corresponding expression values is shown, split by experimental variables, and the DE results for the selected gene, together with automatically generated links to external databases opening up in new tabs, to simplify the subsequent exploration steps.

The content available in the *Gene-Geneset* tab is an excellent starting point to get an overview on the provided data. While navigating the interactive graph, it might occur that the user encounters genes or genesets of particular interest; by simply clicking on the Bookmark button in the header section (or alternatively, pressing the left control key) while the node is selected, these elements are stored throughout the session and collected in the *Bookmarks* panel, where one can generate a dedicated report on these entities.

`GeneTonic` enables the extraction of a graph backbone, wrapping the efficient implementation of the backbone package (Domagalski *et al.*, 2021) to highlight the salient edges of the bipartite projections for each type of features included, as a way to summarize information contained in large networks (Fig. 2F and G).

Additional insight can be extracted by drilling down the interactive *Enrichment Map* (Merico *et al.*, 2010; Yu *et al.*, 2012), either by focusing on the selected nodes (checking out signature heatmaps or bookmarking the genesets for inserting them into the report), or also by running a variety of community detection algorithms on the graph object returned by the `enrichment_map()` function (Suppl. Fig. 1C). Together with the community membership information, it is then possible to obtain a more compact summary for the functional enrichment results, where the most representative genesets for each subpartition of the graph are selected and returned in tabular format. This network-based approach can be exploited to detect the handful of overarching themes, which might give a more immediate snapshot than the many, often redundant, categories, commonly returned by pathway enrichment algorithms (Suppl. Fig. 1E-F-G).

### Summarizing the enrichment results

`GeneTonic` provides numerous ways to summarize the enrichment results, often leveraging the effectiveness of visual representations to extract insights. The *Overview* and *GSViz* panels serve this purpose, showcasing different views on the dataset at hand, with the main controls provided in the right sidebar.

The geneset volcano plot (Fig. 3A) displays all genesets from the `res_enrich` object and labels the most relevant (or any subset of interest). We use one of the aggregated scores (geneset Z-score, or average log_2_ fold change) to determine the horizontal position in the plot. To avoid clutter, it is also possible to reduce the terms based on an overlap threshold, retaining only the most representative ones, and provide this more compact summary to the following visualization routines.

The enhanced table (Fig. 3B) summarizes the top genesets by displaying the log_2_FC of each set’s components along a line (one for each set). On top of the static version, this is provided also as an interactive widget, where tooltips activated with the mouse deliver extra information on each dot, representing a single gene.

The complex relationships among genesets and their behavior across samples are just two aspects one can inspect in depth with the implemented methods. Among these, users can generate a genesets-by-sample heatmap, showing the standardized expression values of the members (via the `gs_scoresheat()` function, Fig. 3C), or alternatively a summary heatmap (with `gs_summary_heat(), Suppl. File 2`), which aims to display the redundancy between different sets, while encoding the values of the expression changes. A multi-dimensional scaling (MDS) plot (Fig. 3D) delivers a 2d visualization of the distance among genesets, based on a similarity measure, e.g. their overlap or other criteria, such as their semantic similarity. In a similar fashion, a dendrogram for genesets enables the possibility to use node color, node size, and branch color to encode relevant features, with the tree structure mirroring the distance matrix based on a similarity measure. `GeneTonic` simplifies the creation of simple summaries for the enrichment, where the essential columns are encoded as graphical parameters of the points, extendable to the case of comparing the same genesets in more than one scenario (e.g. if it is possible to extract more than one contrast from the expression matrix). Switching to polar coordinates, this can be captured in spider plots for one or more `res_enrich` objects (see Suppl. File 2 for more examples of usage).

These visual summaries constitute appealing alternatives to the commonly reported tabular formats, which often fail to provide an overall view for the affected functional landscape.

### Wrapping up the session

The *Bookmarks* panel offers the possibility to review and inspect the shortlisted features of interest, where both genes (on the left side of the interface) and genesets (right side) can be exported to text files.

A more comprehensive report, with dynamically generated content based on the user selections, is compiled when starting the `happy_hour()` function. This is made possible by a template RMarkdown document, included in the `GeneTonic` package, which accesses the input elements and the reactive values for the Shiny components. Notably, this functionality can also be used outside an interactive usage session, specifying as parameters the values for the genes and genesets to focus on. In either case, a full HTML document is rendered, whose content mirrors the structure of the info boxes, and can be later shared or stored as a reproducible artifact for the performed analyses.

Another action button creates the serialized version of a `SummarizedExperiment` object, ready to be provided as the main input to `iSEE` (Rue-Albrecht *et al.*, 2018), for further tailored visualizations, either with standard or custom panels of the web application.

## Discussion

Interpreting the results of transcriptomic studies can be a complex task, where differential expression analysis is combined with a higher-level pathway enrichment analysis, in order to robustly define the molecular actors that display expression changes, and also to identify the underlying functional patterns. Geneset functional enrichment has been successfully applied to thousands of works, and for this step many methods and approaches have been developed. These tasks are also often shared with alternative workflows other than DE analysis, whereas the aim is to extract meaningful information from large lists of genes, yet it is still a prohibitive task to combine in a straightforward way all the single results from each step. This can be for example due to disjoint sets of identifiers, different output and file formats, and to the difficulties in extracting knowledge while handling large numbers of redundant genesets. Providing concise and biologically meaningful views of the underlying cellular processes, defined via differential expression, is essential in many applications, and a proper visualization framework plays a fundamental role in transforming the otherwise tedious and error/bias-prone task of navigating large textual tables into a more compelling activity (Merico *et al.*, 2010; Supek and Škunca, 2017).

In this work, we introduced `GeneTonic` as a solution to explore all the components of a transcriptome dataset in a more integrative way, instead of having to process them as separated outputs. `GeneTonic` is focused on the analytic step devoted to the interpretation of data, rather than on the implementation of additional methods for detection of functionally enriched biological processes or pathways. Consequently, `GeneTonic` implements a variety of summary and visual representations, while accommodating the output of many commonly adopted enrichment tools, making efficient use of the Shiny framework to deliver interactivity and enable drilldown operations. These would otherwise need to be laboriously addressed in multiple iterations of scripted analyses, either done by the user itself or in collaboration with an external unit, such as a bioinformatics core facility.

Several software packages and web-based portals exist for providing similar functionality, and a comprehensive overview of their salient features is presented in Suppl. Table S1. Naturally, these tools differ in terms of implementation, range of applicability, ease of use, with many proposals offering embedded versions of enrichment tests. Since we developed `GeneTonic` in the R programming language, where many such testing procedures are natively available, we instead focused on the support and integration of their output formats into a common workflow. This can be easily combined with existing analysis pipelines, making our tool well suit for potential wide adoption. The comparison with other tools is also available online (https://federicomarini.github.io/GeneTonic_supplement), linked to a Google Sheet where the individual characteristics of each tool can be updated, in order to provide guidance for users who might be seeking advice on which solution best fits their needs (accessible at https://docs.google.com/spreadsheets/d/167XV0w18P0FSld1dt6owN4C2Esxl5FU2QTo4D-wclz0/edit?usp=sharing).

While currently focused on the output of single ORA and FCS enrichment methods, future developments of `GeneTonic` will implement functionality for combined and ensemble approaches, such as `EnrichmentBrowser` (Geistlinger *et al.*, 2016) or `EGSEA` (Alhamdoosh *et al.*, 2016). Moreover, extending such visualizations and interactive summaries to scenarios where multiple omics layers are collected will be a promising avenue for `GeneTonic`, given the growing number of such datasets becoming available. Finally, we intend to address more refined similarity measurements among genesets, e.g. accounting for information contained in protein-protein interaction networks databases (Yoon *et al.*, 2019), in order to better capture the functional relatedness of the affected pathways.

As bioinformatics evolves constantly into a highly interdisciplinary field, it will become increasingly important to develop common platforms usable by many profiles with substantial differences in their level of programming skills, and GeneTonic’s design guidelines adhere to this principle. As a complement to this didactic effect by making the analyses even more open, transparent, and easy to share, `GeneTonic` could make it easier for bioinformatics skilled users to better understand the systems under investigation, prompting e.g. the development of further tailored methods, which could be a key in obtaining a deeper knowledge of the experimental scenarios.

## Conclusion

The identification of relevant functional patterns for the features identified in the differential expression analysis, accounting for the available expression data, remains one of the common bottlenecks for transcriptome-based workflows. `GeneTonic` provides a web application and many underlying functions to assemble the pieces together, supporting the exploration both interactively as well as in a programmatic way. Combining together the results for quantification, DE testing, and functional enrichment (either generated autonomously, or obtained from collaborators), `GeneTonic` assists in the unmet yet increasing need of extracting novel knowledge and insights, which can become daunting especially on larger datasets.

`GeneTonic` has the potential to become an ideal interface between experimental and computational scientists, with the HTML report built via RMarkdown as a milestone for reproducibility, upon conclusion of an interactive session. `GeneTonic` can be integrated in a wide spectrum of existing bioinformatic pipelines, as it provides functions to convert and input the results of many pathway enrichment tools. This aligns with the principle of interoperability at the heart of the Bioconductor project, which enables a large number of such workflows.

The experience of enjoying transcriptomic data analysis and exploration can be easily shared with reduced communication burden, with both experimental and computational sides empowered in the tasks of realizing complex summaries and visualizations. This will significantly facilitate and democratize the discovery process, bridging the gaps existing between technical and domain expertise.

## Additional information

## Acknowledgements

This work has been supported by the computing infrastructure provided by the Core Facility Bioinformatics at the University Medical Center Mainz, used also for deploying the demo instance. The authors thank the members of the Core Facility Bioinformatics at the Institute of Molecular Biology Mainz, Miguel Andrade (IOME, Johannes Gutenberg University of Mainz), Gerrit Toenges and Arsenij Ustjanzew (IMBEI Mainz), and Francesca Finotello (ICBI, Medical University of Innsbruck) for valuable feedback and suggestions.

## Availability and requirements

**Project name:** GeneTonic

**Project home page:** https://bioconductor.org/packages/GeneTonic/ (release), https://github.com/federicomarini/GeneTonic/ (development version)

**Archived version:** https://doi.org/10.5281/zenodo.4742518, package source as gzipped tar archive of the version reported in this article

**Project documentation:** rendered at https://federicomarini.github.io/GeneTonic/

**Operating systems:** Linux, Mac OS, Windows

**Programming language:** R

**Other requirements:** R-4.0.0 or higher, Bioconductor 3.11 or higher

**License:** MIT

**Any restrictions to use by non-academics:** none.

## Abbreviations

DE: Differential Expression
FCS: Functional Class Scoring
FDR: False Discovery Rate
GO: Gene Ontology
GSEA: Gene Set Enrichment Analysis
HGNC: HUGO (Human Genome Organisation) Gene Nomenclature Committee
log_2_FC: base-2 logarithm of the fold change
MA plot: M (log ratio) versus A (mean average) plot
MDS: multi-dimensional scaling
MSigDB: Molecular Signatures Database
NCBI: National Center for Biotechnology Information
ORA: Over-Representation Analysis
PT: Pathway Topology
RNA-seq: RNA sequencing

## Availability of data and materials

The datasets used in this manuscript and its supplements are available from the following articles:

- The data set on the macrophage immune stimulation is included in PubMed ID: 29379200 (https://doi.org/10.1038/s41588-018-0046-7). Dataset deposited at the ENA (ERP020977, project id: PRJEB18997) and accessed from the Bioconductor experiment package macrophage package (https://bioconductor.org/packages/macrophage/, version 1.7.2)
- The data set on murine A20-deficient microglia is included in PubMed ID: 32023471 (https://doi.org/10.1016/j.celrep.2019.12.097). Dataset deposited at the GEO (GSE123033, project id: PRJNA507355) and accessed from the https://github.com/federicomarini/GeneTonic_supplement/repository

The `GeneTonic` package can be downloaded from its Bioconductor page https://bioconductor.org/packages/GeneTonic/ or the GitHub development page https://github.com/federicomarini/GeneTonic/. `GeneTonic` is also provided as a recipe in Bioconda (https://anaconda.org/bioconda/bioconductor-genetonic).

The repository available at https://github.com/federicomarini/GeneTonic_supplement/ contains the code used to generate the supplemental material, and the required input data to replicate the analyses presented in the use cases.

## Authors’ contributions

**Conceptualization:** Federico Marini, Konstantin Strauch

**Data curation:** Federico Marini, Annekathrin Ludt, Jan Linke

**Formal analysis:** Federico Marini, Annekathrin Ludt

**Funding acquisition:** Federico Marini, Konstantin Strauch

**Methodology:** Federico Marini, Annekathrin Ludt

**Project administration:** Federico Marini

**Resources:** Federico Marini, Konstantin Strauch

**Software:** Federico Marini, Annekathrin Ludt, Jan Linke

**Supervision:** Federico Marini, Konstantin Strauch

**Visualization:** Federico Marini, Annekathrin Ludt

**Writing - original draft:** Federico Marini, Konstantin Strauch

**Writing - review & editing:** Federico Marini, Annekathrin Ludt, Jan Linke, Konstantin Strauch All authors read and approved the final version of the manuscript.

## Ethics approval and consent to participate

Not applicable.

## Consent for publication

Not applicable.

---

**Supplemental Figure 1:**
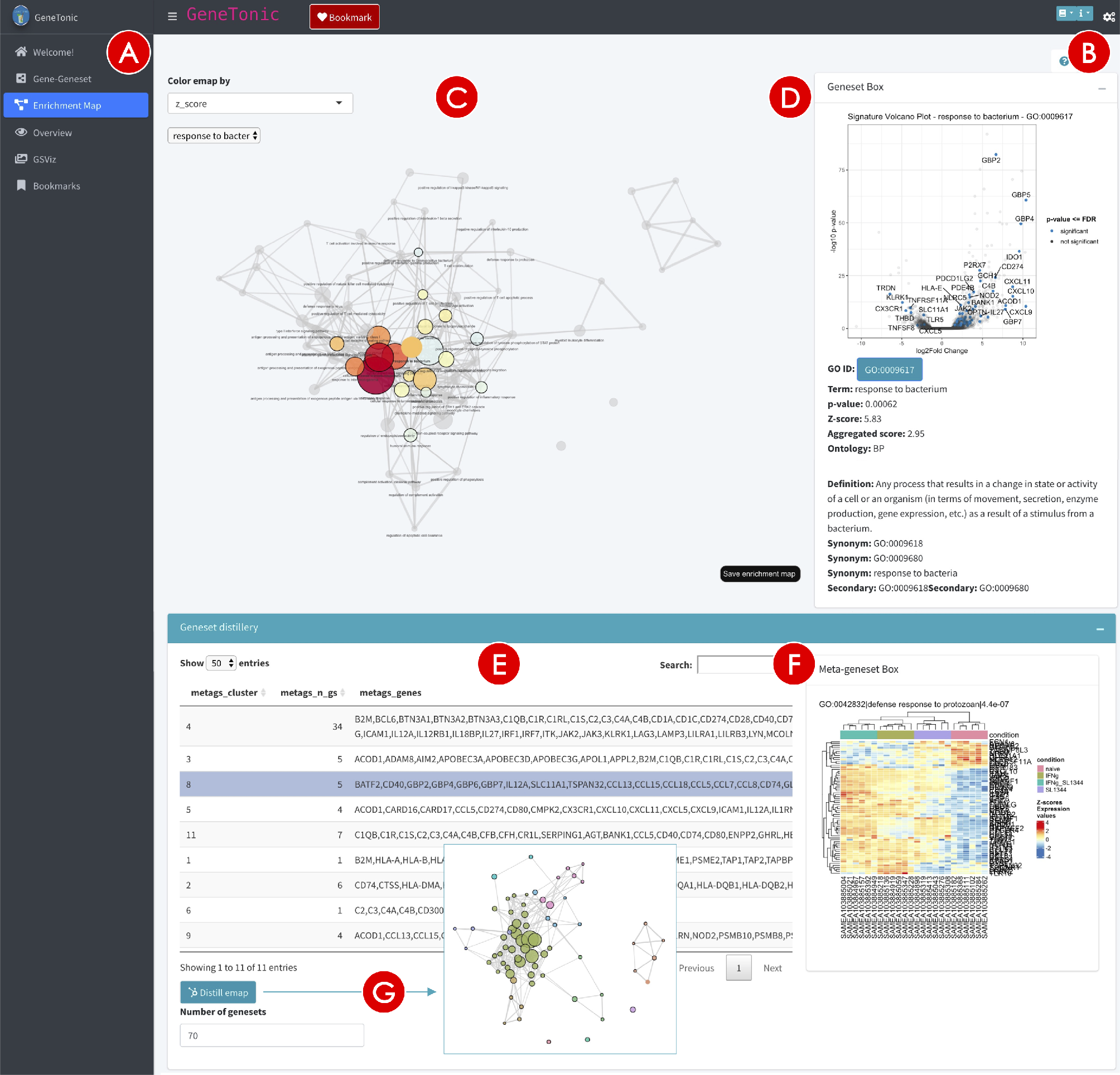
Screenshot of the *Enrichment Map* panel in the GeneTonic application. The sidebar menu (**A**) controls the main navigation in the app, and a common set of options is toggled with the cogs icon (**B**). The main area of the *Enrichment Map* panel (**C**) contains an interactive graph for the enrichment map of the genesets, connected according to their similarity, and color coded according to the specified geneset property (here, the Z-score). Upon clicking on any geneset, a Geneset Box (**D**) is displayed for further exploration (e.g. to show a volcano plot with the geneset members labelled). The geneset distillery (**E**) enables the exploration of meta-genesets, derived by computing clusters on the graph object underlying the enrichment map. From the tabular representation, it is possible to visualize meta-genesets as heatmaps (**F**), or display a modal popup containing the enrichment map where the cluster assignments of the genesets are shown (**G**).

