## Supplemental File 1 for "GeneTonic: an R/Bioconductor package for streamlining the interpretation of RNA-seq data": SupplFile1_usecase_GeneTonic_macrophage.html

Using GeneTonic on the macrophage dataset (ERP020977)


Code 

- Show All Code
- Hide All Code
- Download Rmd

### Using GeneTonic on the macrophage dataset (ERP020977)

###### Federico Marini

Institute of Medical Biostatistics, Epidemiology and Informatics (IMBEI), Mainz  
Center for Thrombosis and Hemostasis (CTH), Mainz  
  
###### Annekathrin Ludt

Institute of Medical Biostatistics, Epidemiology and Informatics (IMBEI), Mainz  
  

###### Jan Linke

Institute of Medical Biostatistics, Epidemiology and Informatics (IMBEI), Mainz  
Center for Thrombosis and Hemostasis (CTH), Mainz  
  

###### Konstantin Strauch

Institute of Medical Biostatistics, Epidemiology and Informatics (IMBEI), Mainz  
  

###### 19 May 2021

### 1 About the data

The data illustrated in this document is an RNA-seq dataset, available at the European Nucleotide Archive under the accession code ERP020977 (https://www.ebi.ac.uk/ena/browser/view/ERP020977).

The data is included in the `macrophage` package available on Bioconductor (https://www.bioconductor.org/packages/release/data/experiment/html/macrophage.html). The data was generated as part of the work to identify shared quantitative trait loci (QTLs) for chromatin accessibility and gene expression in human macrophages (Alasoo et al. 2018) - the manuscript is available at https://www.nature.com/articles/s41588-018-0046-7.

### 2 Loading required packages

We load the packages required to perform all the analytic steps presented in this document.

```
library("DESeq2")
library("topGO")
library("pheatmap")
library("org.Hs.eg.db")
library("pcaExplorer")
library("ideal")
library("DT")
library("GeneTonic")
library("apeglm")
library("dplyr")
library("clusterProfiler")
library("gprofiler2")
library("tibble")
library("fgsea")
library("msigdbr")
library("enrichR")
library("visNetwork")
library("magrittr")
```

### 3 Data processing

After obtaining the data from the `macrophage` package, we generated a `DESeqDataset` object, provided in this repository. The generation of this object is demonstrated in the `GeneTonic` package vignette available through `browseVignette(GeneTonic)` after successful installation of the package. As in the vignette of `GeneTonic`, we will restrict our intention to the comparison between Interferon gamma treated samples vs naive samples of the `macrophage` package.

```
library("macrophage")
library("DESeq2")

data("gse", package = "macrophage")

dds_macrophage <- DESeqDataSet(gse, design = ~line + condition)
# changing the ids to Ensembl instead of the Gencode used in the object
rownames(dds_macrophage) <- substr(rownames(dds_macrophage), 1, 15)
dds_macrophage

# saveRDS(dds_macrophage, file = "usecase_dds_macrophage.rds")
```

#### 3.1 Exploratory data analysis

We read in the dataset, apply the vst transformation for performing PCA and creating a heatmap of the sample to sample distances.

```
dds_macrophage <- readRDS("usecase_dds_macrophage.rds")
dds_macrophage
```

```
## class: DESeqDataSet 
## dim: 58294 24 
## metadata(7): tximetaInfo quantInfo ... txdbInfo version
## assays(3): counts abundance avgTxLength
## rownames(58294): ENSG00000000003 ENSG00000000005 ... ENSG00000285993
##   ENSG00000285994
## rowData names(2): gene_id SYMBOL
## colnames(24): SAMEA103885102 SAMEA103885347 ... SAMEA103885308
##   SAMEA103884949
## colData names(15): names sample_id ... condition line
```

```
vst_macrophage <- vst(dds_macrophage)
vst_macrophage <- vst_macrophage[, vst_macrophage$condition %in% c("naive", "IFNg")]

pheatmap::pheatmap(as.matrix(dist(t(assay(vst_macrophage)))))
```

```
pcaExplorer::pcaplot(vst_macrophage,
                     ntop = 5000,
                     title = "PCA - top5000 most variable genes",
                     ellipse = FALSE
                     )
```

#### 3.2 Differential expression analysis

We set the False Discovery Rate to 0.01 and we run the `DESeq2` workflow, generating results and using the `apeglm` shrinkage estimator.

```
anno_df <- readRDS("usecase_annodf_macrophage.rds")
FDR <- 0.01

dds_macrophage <- DESeq(dds_macrophage)
```

```
results_IFNg_vs_naive <- results(dds_macrophage,
                                 name = "condition_IFNg_vs_naive",
                                 lfcThreshold = 1,
                                 alpha = FDR)
summary(results_IFNg_vs_naive)
```

```
## 
## out of 38829 with nonzero total read count
## adjusted p-value < 0.01
## LFC > 1.00 (up)    : 498, 1.3%
## LFC < -1.00 (down) : 259, 0.67%
## outliers [1]       : 0, 0%
## low counts [2]     : 0, 0%
## (mean count < 0)
## [1] see 'cooksCutoff' argument of ?results
## [2] see 'independentFiltering' argument of ?results
```

```
results_IFNg_vs_naive$SYMBOL <- anno_df$gene_name[match(rownames(results_IFNg_vs_naive), anno_df$gene_id)]

# saveRDS(results_IFNg_vs_naive, file = "usecase_res_de_macrophage.rds")

ideal::plot_ma(results_IFNg_vs_naive, ylim = c(-5, 5), title = "MAplot - IFNg vs Naive")
```

```
tbl_DEres_IFNg_vs_naive <- deseqresult2df(results_IFNg_vs_naive, FDR = FDR)

DT::datatable(tbl_DEres_IFNg_vs_naive, rownames = FALSE)
```

#### 3.3 Functional enrichment analysis

We perform functional enrichment analysis, here using the `topGOtable` wrapper to the method implemented in the `topGO` package.

```
expressedInAssay <- (rowSums(assay(dds_macrophage)) > 0)
geneUniverseExprENS <- rownames(dds_macrophage)[expressedInAssay]
geneUniverseExpr <- anno_df$gene_name[match(geneUniverseExprENS, anno_df$gene_id)]

GObps_IFNg_vs_naive <- topGOtable(
  DEgenes = tbl_DEres_IFNg_vs_naive$SYMBOL,
  BGgenes = geneUniverseExpr,
  ontology = "BP",
  geneID = "symbol",
  addGeneToTerms = TRUE,
  mapping = "org.Hs.eg.db",
  topTablerows = 500
)

GOmfs_IFNg_vs_naive <- topGOtable(
  DEgenes = tbl_DEres_IFNg_vs_naive$SYMBOL,
  BGgenes = geneUniverseExpr,
  ontology = "MF",
  geneID = "symbol",
  addGeneToTerms = TRUE,
  mapping = "org.Hs.eg.db",
  topTablerows = 500
)

GOccs_IFNg_vs_naive <- topGOtable(
  DEgenes = tbl_DEres_IFNg_vs_naive$SYMBOL,
  BGgenes = geneUniverseExpr,
  ontology = "CC",
  geneID = "symbol",
  addGeneToTerms = TRUE,
  mapping = "org.Hs.eg.db",
  topTablerows = 500
)

res_enrich_macrophage <- shake_topGOtableResult(GObps_IFNg_vs_naive)
res_enrich_macrophage <- get_aggrscores(
  res_enrich = res_enrich_macrophage,
  res_de = results_IFNg_vs_naive,
  annotation_obj = anno_df
)

res_enrich <- shake_topGOtableResult(GObps_IFNg_vs_naive)
res_enrich <- get_aggrscores(
  res_enrich = res_enrich,
  res_de = results_IFNg_vs_naive,
  annotation_obj = anno_df
)

# saveRDS(res_enrich, file = "usecase_res_enrich_macrophage.rds")

DT::datatable(res_enrich, rownames = FALSE)
```

##### 3.3.1 Using alternative enrichment analysis methods

It is possible to use the output of different other methods for enrichment analysis, thanks to the `shaker_*` functions implemented in `GeneTonic` - see for example `?shake_topGOtableResult`, and navigate to the manual pages for the other functions listed in the “See Also” section.

`GeneTonic` is able to convert for you the output of

- DAVID (text file, downloaded from the DAVID website)
- clusterProfiler (takes `enrichResult` objects)
- enrichr (a data.frame with the output of `enrichr`, or the text file exported from Enrichr)
- fgsea (the output of `fgsea()` function)
- g:Profiler (the text file output as exported from g:Profiler, or a data.frame with the output of `gost()` in `gprofiler2`)

Some examples on how to use them are reported here:

```
# clusterProfiler --------------------------------------------------------------
degenes_macrophage <- deseqresult2df(results_IFNg_vs_naive, FDR = FDR)$id
ego_macrophage <- enrichGO(gene          = degenes_macrophage,
                           universe      = rownames(dds_macrophage)[expressedInAssay],
                           OrgDb         = org.Hs.eg.db,
                           ont           = "BP",
                           keyType       = 'ENSEMBL',
                           pAdjustMethod = "BH",
                           pvalueCutoff  = 0.01,
                           qvalueCutoff  = 0.05,
                           readable      = TRUE)
res_enrich_clusterprofiler <- shake_enrichResult(ego_macrophage)

# g:profiler -------------------------------------------------------------------
degenes <- deseqresult2df(results_IFNg_vs_naive, FDR = FDR)$SYMBOL
gostres_macrophage <- gost(
  query = degenes, 
  ordered_query = FALSE, 
  multi_query = FALSE, 
  significant = FALSE, 
  exclude_iea = TRUE, 
  measure_underrepresentation = FALSE, 
  evcodes = TRUE, 
  user_threshold = 0.05, 
  correction_method = "g_SCS", 
  domain_scope = "annotated", 
  numeric_ns = "", 
  sources = "GO:BP", 
  as_short_link = FALSE)

res_enrich_gprofiler <- shake_gprofilerResult(gprofiler_output = gostres_macrophage$result)

# fgsea ------------------------------------------------------------------------
res2 <- results_IFNg_vs_naive %>%
  as.data.frame() %>% 
  dplyr::select(SYMBOL, log2FoldChange)
de_ranks <- deframe(res2)
msigdbr_df <- msigdbr(species = "Homo sapiens", category = "C5", subcategory = "BP")
msigdbr_list <- split(x = msigdbr_df$gene_symbol, f = msigdbr_df$gs_name)
fgseaRes <- fgsea(pathways = msigdbr_list, 
                  stats = de_ranks, 
                  nperm=100000)
fgseaRes <- fgseaRes %>% 
  arrange(desc(NES))

res_enrich_fgsea <- shake_fgseaResult(fgsea_output = fgseaRes)

# enrichr ----------------------------------------------------------------------
dbs <- c("GO_Molecular_Function_2018",
         "GO_Cellular_Component_2018",
         "GO_Biological_Process_2018",
         "KEGG_2019_Human",
         "Reactome_2016",
         "WikiPathways_2019_Human")
degenes <- deseqresult2df(results_IFNg_vs_naive, FDR = FDR)$SYMBOL
enrichr_output_macrophage <- enrichr(degenes, dbs)

res_enrich_enrichr_BPs <- shake_enrichrResult(
  enrichr_output = enrichr_output_macrophage$GO_Biological_Process_2018)
res_enrich_enrichr_KEGG <- shake_enrichrResult(
  enrichr_output = enrichr_output_macrophage$KEGG_2019_Human)
```

#### 3.4 Assembling the `gtl` object

To simplify the usage of the function from `GeneTonic`, we create a `GeneTonic_list` object.

```
gtl_macrophage <- GeneTonic_list(
  dds = dds_macrophage,
  res_de = results_IFNg_vs_naive,
  res_enrich = res_enrich,
  annotation_obj = anno_df
)

# saveRDS(gtl_macrophage, "gtl_macrophage.rds")
```

### 4 Running `GeneTonic` on the dataset

Let’s load the `GeneTonicList` object, containing all the structured input needed.

```
gtl_macrophage <- readRDS("gtl_macrophage.rds")
```

#### 4.1 `GeneTonic`, interactively

This command will launch the `GeneTonic` app:

```
GeneTonic(gtl = gtl_macrophage)
```

The exploration can be guided by launching the introductory tours for each section of the app, and finally one can generate a small report focused on the features and genesets of interest.

If some more custom visualizations are required, it is possible to export the objects in use from the app as a `SummarizedExperiment` object, to be then passed to the `iSEE` software (Rue-Albrecht et al. 2018) for further exploration.

#### 4.2 Using `GeneTonic`’s functions in analysis reports

The functionality of `GeneTonic` can be used also as standalone functions, to be called for example in existing analysis reports in RMarkdown, or R scripts.

```
em <- enrichment_map(gtl = gtl_macrophage, 
                     n_gs = 100,
                     color_by = "z_score")

em %>%
  visIgraph() %>%
  visOptions(
    highlightNearest = list(
      enabled = TRUE,
      degree = 1,
      hover = TRUE
    ),
    nodesIdSelection = TRUE
  )
```

Focusing on individual genesets, we can use `gs_heatmap()` and `gs_volcano()` as in the lines that follow:

```
gs_heatmap(
  se = vst_macrophage,
  gtl = gtl_macrophage,
  geneset_id = res_enrich$gs_id[1],
  cluster_columns = TRUE,
  FDR = FDR)
```

```
signature_volcano(
  gtl = gtl_macrophage,
  geneset_id = res_enrich$gs_id[1],
  FDR = FDR
)
```

If an overview of the affected pathways is desired, one can apply the fuzzy clustering algorithm on the enrichment results, and obtain visual summaries focused on the genesets identified as representative ones of each cluster.

```
res_enrich_subset <- res_enrich[1:100, ]
fuzzy_subset <- gs_fuzzyclustering(
  res_enrich = res_enrich_subset,
  n_gs = nrow(res_enrich_subset),
  gs_ids = NULL,
  similarity_matrix = NULL,
  similarity_threshold = 0.35,
  fuzzy_seeding_initial_neighbors = 3,
  fuzzy_multilinkage_rule = 0.5
)

# show all genesets members of the first cluster
DT::datatable(
  fuzzy_subset[fuzzy_subset$gs_fuzzycluster == "1", ]
)
```

```
# list only the representative clusters
DT::datatable(
  fuzzy_subset[fuzzy_subset$gs_cluster_status == "Representative", ]
)
```

Let’s focus on the top 15 sets identified with the fuzzy clustering.

```
focus_genesets <- 
  head(fuzzy_subset$gs_id[fuzzy_subset$gs_cluster_status == "Representative"], 15)

score_mat <- gs_scores(se = vst_macrophage,
                       gtl = gtl_macrophage)

gs_scoresheat(score_mat,
              n_gs = 0,
              gs_ids = focus_genesets)
```

```
gs_mds(
  gtl = gtl_macrophage,
  n_gs = 200,
  mds_labels = 0,
  gs_labels = focus_genesets)
```

More examples on the functions to use and their applications is also included in the other supplemental file, on the A20-deletion in microglia (Mohebiany et al. 2020).

### Session information

```
sessionInfo()
```

```
## R version 4.1.0 RC (2021-05-17 r80314)
## Platform: x86_64-apple-darwin17.0 (64-bit)
## Running under: macOS Mojave 10.14.6
## 
## Matrix products: default
## BLAS:   /System/Library/Frameworks/Accelerate.framework/Versions/A/Frameworks/vecLib.framework/Versions/A/libBLAS.dylib
## LAPACK: /Library/Frameworks/R.framework/Versions/4.1/Resources/lib/libRlapack.dylib
## 
## locale:
## [1] en_US.UTF-8/en_US.UTF-8/en_US.UTF-8/C/en_US.UTF-8/en_US.UTF-8
## 
## attached base packages:
## [1] parallel  stats4    stats     graphics  grDevices utils     datasets 
## [8] methods   base     
## 
## other attached packages:
##  [1] edgeR_3.33.7                limma_3.47.15              
##  [3] DEFormats_1.19.0            igraph_1.2.6               
##  [5] org.Mm.eg.db_3.13.0         magrittr_2.0.1             
##  [7] visNetwork_2.0.9            enrichR_3.0                
##  [9] msigdbr_7.4.1               fgsea_1.17.0               
## [11] tibble_3.1.2                gprofiler2_0.2.0           
## [13] clusterProfiler_3.99.2      dplyr_1.0.6                
## [15] apeglm_1.13.4               GeneTonic_1.3.8            
## [17] DT_0.18                     ideal_1.15.2               
## [19] pcaExplorer_2.17.1          bigmemory_4.5.36           
## [21] org.Hs.eg.db_3.13.0         pheatmap_1.0.12            
## [23] topGO_2.43.0                SparseM_1.81               
## [25] GO.db_3.13.0                AnnotationDbi_1.53.2       
## [27] graph_1.69.0                DESeq2_1.31.21             
## [29] SummarizedExperiment_1.21.3 Biobase_2.51.0             
## [31] MatrixGenerics_1.3.1        matrixStats_0.58.0         
## [33] GenomicRanges_1.43.4        GenomeInfoDb_1.27.13       
## [35] IRanges_2.25.11             S4Vectors_0.29.19          
## [37] BiocGenerics_0.37.6         knitr_1.33                 
## [39] job_0.2                     rmarkdown_2.8              
## 
## loaded via a namespace (and not attached):
##   [1] ps_1.6.0                 Rsamtools_2.7.2          foreach_1.5.1           
##   [4] rprojroot_2.0.2          crayon_1.4.1             MASS_7.3-54             
##   [7] backports_1.2.1          nlme_3.1-152             colourpicker_1.1.0      
##  [10] GOSemSim_2.17.2          rlang_0.4.11             XVector_0.31.1          
##  [13] callr_3.7.0              filelock_1.0.2           GOstats_2.57.0          
##  [16] BiocParallel_1.25.5      rjson_0.2.20             bit64_4.0.5             
##  [19] glue_1.4.2               rngtools_1.5             processx_3.5.2          
##  [22] UpSetR_1.4.0             shinyAce_0.4.1           shinydashboard_0.7.1    
##  [25] DOSE_3.17.2              tidyselect_1.1.1         XML_3.99-0.6            
##  [28] tidyr_1.1.3              GenomicAlignments_1.27.2 xtable_1.8-4            
##  [31] evaluate_0.14            ggplot2_3.3.3            cli_2.5.0               
##  [34] zlibbioc_1.37.0          rstudioapi_0.13          miniUI_0.1.1.1          
##  [37] bslib_0.2.5              fastmatch_1.1-0          BiocStyle_2.19.2        
##  [40] treeio_1.15.7            shiny_1.6.0              xfun_0.23               
##  [43] clue_0.3-59              pkgbuild_1.2.0           cluster_2.1.2           
##  [46] caTools_1.18.2           tidygraph_1.2.0          TSP_1.1-10              
##  [49] KEGGREST_1.31.2          expm_0.999-6             ggrepel_0.9.1           
##  [52] threejs_0.3.3            ape_5.5                  dendextend_1.15.1       
##  [55] shinyWidgets_0.6.0       Biostrings_2.59.4        png_0.1-7               
##  [58] withr_2.4.2              shinyBS_0.61             bitops_1.0-7            
##  [61] slam_0.1-48              ggforce_0.3.3            RBGL_1.67.0             
##  [64] plyr_1.8.6               GSEABase_1.53.2          coda_0.19-4             
##  [67] pillar_1.6.1             gplots_3.1.1             GlobalOptions_0.1.2     
##  [70] cachem_1.0.5             GenomicFeatures_1.43.8   GetoptLong_1.0.5        
##  [73] vctrs_0.3.8              ellipsis_0.3.2           generics_0.1.0          
##  [76] NMF_0.23.0               tools_4.1.0              munsell_0.5.0           
##  [79] tweenr_1.0.2             DelayedArray_0.17.13     fastmap_1.1.0           
##  [82] compiler_4.1.0           httpuv_1.6.1             rtracklayer_1.51.5      
##  [85] geneLenDataBase_1.27.0   pkgmaker_0.32.2          plotly_4.9.3            
##  [88] GenomeInfoDbData_1.2.6   gridExtra_2.3            lattice_0.20-44         
##  [91] AnnotationForge_1.33.6   utf8_1.2.1               later_1.2.0             
##  [94] BiocFileCache_1.99.8     jsonlite_1.7.2           scales_1.1.1            
##  [97] tidytree_0.3.3           genefilter_1.73.1        lazyeval_0.2.2          
## [100] promises_1.2.0.1         doParallel_1.0.16        checkmate_2.0.0         
## [103] goseq_1.43.0             cowplot_1.1.1            webshot_0.5.2           
## [106] downloader_0.4           survival_3.2-11          numDeriv_2016.8-1.1     
## [109] rsconnect_0.8.17         yaml_2.2.1               rintrojs_0.3.0          
## [112] htmltools_0.5.1.1        memoise_2.0.0            spelling_2.2            
## [115] BiocIO_1.1.2             locfit_1.5-9.4           seriation_1.2-9         
## [118] graphlayouts_0.7.1       viridisLite_0.4.0        digest_0.6.27           
## [121] assertthat_0.2.1         mime_0.10                commonmark_1.7          
## [124] rappdirs_0.3.3           emdbook_1.3.12           registry_0.5-1          
## [127] BiasedUrn_1.07           bigmemory.sri_0.1.3      RSQLite_2.2.7           
## [130] remotes_2.3.0            data.table_1.14.0        blob_1.2.1              
## [133] lpsymphony_1.19.0        labeling_0.4.2           splines_4.1.0           
## [136] Cairo_1.5-12.2           RCurl_1.98-1.3           hms_1.1.0               
## [139] colorspace_2.0-1         base64enc_0.1-3          BiocManager_1.30.15     
## [142] shape_1.4.5              aplot_0.0.6              sass_0.4.0              
## [145] Rcpp_1.0.6               bookdown_0.22            mvtnorm_1.1-1           
## [148] circlize_0.4.12          enrichplot_1.11.3        backbone_1.4.0          
## [151] fansi_0.4.2              IHW_1.19.0               R6_2.5.0                
## [154] grid_4.1.0               lifecycle_1.0.0          curl_4.3.1              
## [157] jquerylib_0.1.4          DO.db_2.9                Matrix_1.3-3            
## [160] qvalue_2.23.0            RColorBrewer_1.1-2       iterators_1.0.13        
## [163] stringr_1.4.0            htmlwidgets_1.5.3        hunspell_3.0.1          
## [166] polyclip_1.10-0          biomaRt_2.47.9           purrr_0.3.4             
## [169] crosstalk_1.1.1          shadowtext_0.0.8         ComplexHeatmap_2.7.11   
## [172] mgcv_1.8-35              patchwork_1.1.1          bdsmatrix_1.3-4         
## [175] codetools_0.2-18         bs4Dash_0.5.0            gtools_3.8.2            
## [178] prettyunits_1.1.1        dbplyr_2.1.1             gridBase_0.4-7          
## [181] gtable_0.3.0             DBI_1.1.1                dynamicTreeCut_1.63-1   
## [184] highr_0.9                httr_1.4.2               KernSmooth_2.23-20      
## [187] stringi_1.6.2            progress_1.2.2           reshape2_1.4.4          
## [190] farver_2.1.0             heatmaply_1.2.1          annotate_1.69.2         
## [193] viridis_0.6.1            rentrez_1.2.3            fdrtool_1.2.16          
## [196] Rgraphviz_2.35.0         magick_2.7.2             ggtree_2.99.0           
## [199] xml2_1.3.2               rvcheck_0.1.8            bbmle_1.0.23.1          
## [202] restfulr_0.0.13          geneplotter_1.69.0       Category_2.57.2         
## [205] bit_4.0.4                shinycssloaders_1.0.0    scatterpie_0.1.6        
## [208] ggraph_2.0.5             babelgene_21.4           pkgconfig_2.0.3
```
