## Supplemental File 2 for "GeneTonic: an R/Bioconductor package for streamlining the interpretation of RNA-seq data": SupplFile2_usecase_GeneTonic_microgliaA20.html

Using GeneTonic on the A20-deficient microglia dataset (GSE123033)


Code 

- Show All Code
- Hide All Code
- Download Rmd

### Using GeneTonic on the A20-deficient microglia dataset (GSE123033)

###### Federico Marini

Institute of Medical Biostatistics, Epidemiology and Informatics (IMBEI), Mainz  
Center for Thrombosis and Hemostasis (CTH), Mainz  
  
###### Annekathrin Ludt

Institute of Medical Biostatistics, Epidemiology and Informatics (IMBEI), Mainz  
  

###### Jan Linke

Institute of Medical Biostatistics, Epidemiology and Informatics (IMBEI), Mainz  
Center for Thrombosis and Hemostasis (CTH), Mainz  
  

###### Konstantin Strauch

Institute of Medical Biostatistics, Epidemiology and Informatics (IMBEI), Mainz  
  

###### 19 May 2021

### 1 About the data

The data illustrated in this document is an RNA-seq dataset, available at the GEO repository under the accession code GSE123033 (https://www.ncbi.nlm.nih.gov/geo/query/acc.cgi?acc=GSE123033).

### 2 Loading required packages

We load the packages required to perform all the analytic steps presented in this document.

```
library("DESeq2")
library("topGO")
library("pheatmap")
library("org.Mm.eg.db")
library("pcaExplorer")
library("ideal")
library("DT")
library("GeneTonic")
```

### 3 Data processing

After obtaining a count matrix (via STAR and featureCounts), we generated a `DESeqDataset` object, provided in this repository for the ease of reproducing the presented analyses.

#### 3.1 Exploratory data analysis

We read in the dataset, apply the rlog transformation for performing PCA and creating a heatmap of the sample to sample distances.

```
dds_alma <- readRDS("usecase_dds_alma.rds")
dds_alma
```

```
## class: DESeqDataSet 
## dim: 41388 5 
## metadata(1): version
## assays(4): counts mu H cooks
## rownames(41388): ENSMUSG00000090025 ENSMUSG00000064842 ...
##   ENSMUSG00000096730 ENSMUSG00000095742
## rowData names(22): baseMean baseVar ... deviance maxCooks
## colnames(5): control_R1 control_R2 ko_del_R1 ko_del_R2 ko_del_R3
## colData names(8): fastqfile bamfile ... sampleName sizeFactor
```

```
rld_alma <- rlogTransformation(dds_alma)

anno_df <- readRDS("usecase_annodf_alma.rds")

pheatmap::pheatmap(as.matrix(dist(t(assay(rld_alma)))))
```

```
pcaExplorer::pcaplot(rld_alma,
  ntop = 5000,
  title = "PCA - top5000 most variable genes",
  ellipse = FALSE
)
```

#### 3.2 Differential expression analysis

We set the False Discovery Rate to 0.05 and we run the `DESeq2` workflow, generating results and using the `apeglm` shrinkage estimator.

```
FDR <- 0.05

dds_alma <- DESeq(dds_alma)

res_alma_ko_vs_ctrl <- results(dds_alma, name = "condition_ko_del_vs_control", alpha = FDR)
summary(res_alma_ko_vs_ctrl)
```

```
## 
## out of 20699 with nonzero total read count
## adjusted p-value < 0.05
## LFC > 0 (up)       : 668, 3.2%
## LFC < 0 (down)     : 342, 1.7%
## outliers [1]       : 3, 0.014%
## low counts [2]     : 7984, 39%
## (mean count < 5)
## [1] see 'cooksCutoff' argument of ?results
## [2] see 'independentFiltering' argument of ?results
```

```
library("apeglm")
res_alma_ko_vs_ctrl <- lfcShrink(dds_alma, coef = "condition_ko_del_vs_control", type = "apeglm", res = res_alma_ko_vs_ctrl)

res_alma_ko_vs_ctrl$SYMBOL <- anno_df$gene_name[match(rownames(res_alma_ko_vs_ctrl), anno_df$gene_id)]

summary(res_alma_ko_vs_ctrl)
```

```
## 
## out of 20699 with nonzero total read count
## adjusted p-value < 0.05
## LFC > 0 (up)       : 668, 3.2%
## LFC < 0 (down)     : 342, 1.7%
## outliers [1]       : 3, 0.014%
## low counts [2]     : 7984, 39%
## (mean count < 5)
## [1] see 'cooksCutoff' argument of ?results
## [2] see 'independentFiltering' argument of ?results
```

```
# saveRDS(res_alma_ko_vs_ctrl, file = "usecase_res_de_alma.rds")

ideal::plot_ma(res_alma_ko_vs_ctrl, ylim = c(-5, 5), title = "MAplot - A20ko vs Ctrl")
```

```
tbl_DEres_alma_ko_vs_ctrl <- deseqresult2df(res_alma_ko_vs_ctrl, FDR = FDR)

DT::datatable(tbl_DEres_alma_ko_vs_ctrl, rownames = FALSE)
```

#### 3.3 Functional enrichment analysis

We perform functional enrichment analysis, here using the `topGOtable` wrapper to the method implemented in the `topGO` package.

```
expressedInAssay <- (rowSums(assay(dds_alma)) > 0)
geneUniverseExprENS <- rownames(dds_alma)[expressedInAssay]
geneUniverseExpr <- anno_df$gene_name[match(geneUniverseExprENS, anno_df$gene_id)]

GObps_ko_vs_ctrl <- topGOtable(
  DEgenes = tbl_DEres_alma_ko_vs_ctrl$SYMBOL,
  BGgenes = geneUniverseExpr,
  ontology = "BP",
  geneID = "symbol",
  addGeneToTerms = TRUE,
  mapping = "org.Mm.eg.db",
  topTablerows = 500
)

GOmfs_ko_vs_ctrl <- topGOtable(
  DEgenes = tbl_DEres_alma_ko_vs_ctrl$SYMBOL,
  BGgenes = geneUniverseExpr,
  ontology = "MF",
  geneID = "symbol",
  addGeneToTerms = TRUE,
  mapping = "org.Mm.eg.db",
  topTablerows = 500
)

GOccs_ko_vs_ctrl <- topGOtable(
  DEgenes = tbl_DEres_alma_ko_vs_ctrl$SYMBOL,
  BGgenes = geneUniverseExpr,
  ontology = "CC",
  geneID = "symbol",
  addGeneToTerms = TRUE,
  mapping = "org.Mm.eg.db",
  topTablerows = 500
)

res_enrich_a20 <- shake_topGOtableResult(GObps_ko_vs_ctrl)
res_enrich_a20 <- get_aggrscores(
  res_enrich = res_enrich_a20,
  res_de = res_alma_ko_vs_ctrl,
  annotation_obj = anno_df
)

# saveRDS(res_enrich_a20, file = "usecase_res_enrich_alma.rds")

DT::datatable(res_enrich_a20, rownames = FALSE)
```

##### 3.3.1 Using alternative enrichment analysis methods

It is possible to use the output of different other methods for enrichment analysis, thanks to the `shaker_*` functions implemented in `GeneTonic` - see for example `?shake_topGOtableResult`, and navigate to the manual pages for the other functions listed in the “See Also” section.

`GeneTonic` is able to convert for you the output of

- DAVID (text file, downloaded from the DAVID website)
- clusterProfiler (takes `enrichResult` objects)
- enrichr (a data.frame with the output of `enrichr`, or the text file exported from Enrichr)
- fgsea (the output of `fgsea()` function)
- g:Profiler (the text file output as exported from g:Profiler, or a data.frame with the output of `gost()` in `gprofiler2`)

Some examples on how to use them are reported here:

```
# clusterProfiler --------------------------------------------------------------
library("clusterProfiler")
degenes_alma <- deseqresult2df(res_alma_ko_vs_ctrl, FDR = 0.05)$id
ego_alma <- enrichGO(
  gene = degenes_alma,
  universe = rownames(dds_alma)[expressedInAssay],
  OrgDb = org.Mm.eg.db,
  ont = "BP",
  keyType = "ENSEMBL",
  pAdjustMethod = "BH",
  pvalueCutoff = 0.01,
  qvalueCutoff = 0.05,
  readable = TRUE
)
res_enrich_clusterprofiler <- shake_enrichResult(ego_alma)

# g:profiler -------------------------------------------------------------------
library("gprofiler2")
degenes <- deseqresult2df(res_alma_ko_vs_ctrl, FDR = 0.05)$SYMBOL
gostres_a20 <- gost(
  query = degenes,
  ordered_query = FALSE,
  multi_query = FALSE,
  significant = FALSE,
  exclude_iea = TRUE,
  measure_underrepresentation = FALSE,
  evcodes = TRUE,
  user_threshold = 0.05,
  correction_method = "g_SCS",
  domain_scope = "annotated",
  numeric_ns = "",
  sources = "GO:BP",
  as_short_link = FALSE
)

res_enrich_gprofiler <- shake_gprofilerResult(gprofiler_output = gostres_a20$result)

# fgsea ------------------------------------------------------------------------
library("dplyr")
library("tibble")
library("fgsea")
library("msigdbr")
res2 <- res_alma_ko_vs_ctrl %>%
  as.data.frame() %>%
  dplyr::select(SYMBOL, log2FoldChange)
de_ranks <- deframe(res2)
msigdbr_df <- msigdbr(species = "Mus musculus", category = "C5", subcategory = "BP")
msigdbr_list <- split(x = msigdbr_df$gene_symbol, f = msigdbr_df$gs_name)
fgseaRes <- fgsea(
  pathways = msigdbr_list,
  stats = de_ranks,
  nperm = 100000
)
fgseaRes <- fgseaRes %>%
  arrange(desc(NES))

res_enrich_fgsea <- shake_fgseaResult(fgsea_output = fgseaRes)

# enrichr ----------------------------------------------------------------------
library("enrichR")
dbs <- c(
  "GO_Molecular_Function_2018",
  "GO_Cellular_Component_2018",
  "GO_Biological_Process_2018",
  "KEGG_2019_Human",
  "Reactome_2016",
  "WikiPathways_2019_Human"
)
degenes <- (deseqresult2df(res_alma_ko_vs_ctrl, FDR = 0.05)$SYMBOL)
enrichr_output_a20 <- enrichr(degenes, dbs)

res_enrich_enrichr_BPs <- shake_enrichrResult(
  enrichr_output = enrichr_output_a20$GO_Biological_Process_2018
)
res_enrich_enrichr_KEGG <- shake_enrichrResult(
  enrichr_output = enrichr_output_a20$KEGG_2019_Human
)
```

#### 3.4 Assembling the `gtl` object

To simplify the usage of the function from `GeneTonic`, we create a `GeneTonic_list` object

```
gtl_alma <- GeneTonic_list(
  dds = dds_alma,
  res_de = res_alma_ko_vs_ctrl,
  res_enrich = res_enrich_a20,
  annotation_obj = anno_df
)

# saveRDS(gtl_alma, "gtl_alma.rds")
```

### 4 Running `GeneTonic` on the dataset

Let’s load the `GeneTonicList` object, containing all the structured input needed.

```
gtl_alma <- readRDS("gtl_alma.rds")
```

#### 4.1 `GeneTonic`, interactively

This command will launch the `GeneTonic` app:

```
GeneTonic(gtl = gtl_alma)
```

The exploration can be guided by launching the introductory tours for each section of the app, and finally one can generate a small report focused on the features and genesets of interest.

If some more custom visualizations are required, it is possible to export the objects in use from the app as a SummarizedExperiment object, to be then passed to the `iSEE` software (Rue-Albrecht et al. 2018) for further exploration.

#### 4.2 Using `GeneTonic`’s functions in analysis reports

The functionality of `GeneTonic` can be used also as standalone functions, to be called for example in existing analysis reports in RMarkdown, or R scripts.

For more details on the functions and their outputs, please refer to the package documentation and the vignette.

##### 4.2.1 Genesets and genes

We can represent the relationships between genesets and genes with `enhance_table()`, `gs_summary_heat()`, and `ggs_graph()`:

```
p <- enhance_table(
  gtl = gtl_alma,
  n_gs = 30,
  chars_limit = 60
)
p
```

```
## could be made interactive with...
# library("plotly")
# ggplotly(p)

gs_summary_heat(
  gtl = gtl_alma,
  n_gs = 5
)
```

```
ggs <- ggs_graph(
  gtl = gtl_alma,
  n_gs = 20
)
ggs
```

```
## IGRAPH 5307862 UN-- 291 626 -- 
## + attr: name (v/c), nodetype (v/c), shape (v/c), color (v/c), title
## | (v/c)
## + edges from 5307862 (vertex names):
##  [1] cellular response to interferon-beta--Gbp2   
##  [2] cellular response to interferon-beta--Gbp3   
##  [3] cellular response to interferon-beta--Gbp6   
##  [4] cellular response to interferon-beta--Gm12185
##  [5] cellular response to interferon-beta--Gm4841 
##  [6] cellular response to interferon-beta--Gm4951 
##  [7] cellular response to interferon-beta--Ifi204 
## + ... omitted several edges
```

```
## could be viewed interactively with
library("visNetwork")
library("magrittr")
ggs %>%
  visIgraph() %>%
  visOptions(
    highlightNearest = list(
      enabled = TRUE,
      degree = 1,
      hover = TRUE
    ),
    nodesIdSelection = TRUE
  )
```

##### 4.2.2 Overview representations

We can also obtain summary representations of the enrichment results, either as table-like views (with `gs_volcano()`, `gs_summary_overview()`, `gs_fuzzyclustering()`, `gs_scoresheat()`), or to represent the relationships among genesets (via `enrichment_map()`, `gs_dendro()`, `gs_mds()`)

```
gs_volcano(
  res_enrich = res_enrich_a20,
  p_threshold = 0.05,
  color_by = "aggr_score",
  volcano_labels = 5,
  gs_ids = c("GO:0033690", "GO:0050770"),
  plot_title = "A20-microglia geneset volcano"
)
```

```
gs_summary_overview(
  res_enrich = res_enrich_a20,
  n_gs = 60,
  p_value_column = "gs_pvalue",
  color_by = "z_score"
)
```

```
res_enrich_subset <- res_enrich_a20[1:200, ]
fuzzy_subset <- gs_fuzzyclustering(
  res_enrich = res_enrich_subset,
  n_gs = nrow(res_enrich_subset),
  gs_ids = NULL,
  similarity_matrix = NULL,
  similarity_threshold = 0.35,
  fuzzy_seeding_initial_neighbors = 3,
  fuzzy_multilinkage_rule = 0.5
)

# show all genesets members of the first cluster
DT::datatable(
  fuzzy_subset[fuzzy_subset$gs_fuzzycluster == "1", ]
)
```

```
# list only the representative clusters
DT::datatable(
  fuzzy_subset[fuzzy_subset$gs_cluster_status == "Representative", ]
)
```

```
vst_alma <- vst(dds_alma)
scores_mat <- gs_scores(
  se = vst_alma,
  gtl = gtl_alma
)
gs_scoresheat(scores_mat,
  n_gs = 30
)
```

```
em <- enrichment_map(gtl = gtl_alma, n_gs = 100)

library("visNetwork")
library("magrittr")
em %>%
  visIgraph() %>%
  visOptions(
    highlightNearest = list(
      enabled = TRUE,
      degree = 1,
      hover = TRUE
    ),
    nodesIdSelection = TRUE
  )
```

```
distilled <- distill_enrichment(
  gtl = gtl_alma,
  n_gs = 100,
  cluster_fun = "cluster_markov"
)
DT::datatable(distilled$distilled_table)
```

```
DT::datatable(distilled$res_enrich)
```

```
dg <- distilled$distilled_em

library("igraph")
library("visNetwork")
library("magrittr")

# defining a color palette for nicer display
colpal <- colorspace::rainbow_hcl(length(unique(V(dg)$color)))[V(dg)$color]
V(dg)$color.background <- scales::alpha(colpal, alpha = 0.8)
V(dg)$color.highlight <- scales::alpha(colpal, alpha = 1)
V(dg)$color.hover <- scales::alpha(colpal, alpha = 0.5)

V(dg)$color.border <- "black"

visNetwork::visIgraph(dg) %>%
  visOptions(
    highlightNearest = list(
      enabled = TRUE,
      degree = 1,
      hover = TRUE
    ),
    nodesIdSelection = TRUE,
    selectedBy = "membership"
  )
```

```
gs_dendro(
  gtl = gtl_alma,
  n_gs = 50,
  gs_dist_type = "kappa",
  clust_method = "ward.D2",
  color_leaves_by = "z_score",
  size_leaves_by = "gs_pvalue",
  color_branches_by = "clusters",
  create_plot = TRUE
)
```

```
gs_mds(
  gtl = gtl_alma,
  n_gs = 200,
  gs_ids = NULL,
  similarity_measure = "kappa_matrix",
  mds_k = 2,
  mds_labels = 10,
  mds_colorby = "z_score",
  gs_labels = NULL,
  plot_title = NULL
)
```

##### 4.2.3 General use functions

It is possible to obtain focused representations and descriptions of single genesets with `gs_heatmap()`, `go2html()`, and `signature_volcano()` - these need the Gene Ontology identifier to generate the desired content, while the `gtl` object contains all the necessary expression matrices and tables.

```
gs_heatmap(
  se = vst_alma,
  gtl = gtl_alma,
  geneset_id = "GO:0060337",
  cluster_columns = TRUE,
  anno_col_info = "condition"
)
```

```
go_2_html("GO:0060337",
  res_enrich = res_enrich_a20
)
```

**GO ID:** GO:0060337

```
signature_volcano(
  gtl = gtl_alma,
  geneset_id = "GO:0060337",
  FDR = 0.05,
  color = "#1a81c2"
)
```

As an example, these plots are related to the data presented in Figure 3 of (Mohebiany et al. 2020).

```
genes_synapse_assembly <- c(
  "Bhlhb9",
  "Mef2c",
  "Clstn1",
  "Amigo1",
  "Eef2k",
  "Ephb3",
  "Thbs2",
  "Pdlim5"
)

genes_synapse_assembly_ensids <-
  anno_df$gene_id[match(genes_synapse_assembly, anno_df$gene_name)]

gs_heatmap(
  se = vst_alma,
  gtl = gtl_alma,
  genelist = genes_synapse_assembly_ensids,
  cluster_columns = TRUE,
  cluster_rows = FALSE,
  anno_col_info = "condition"
)
```

```
res_enrich_a20[res_enrich_a20$gs_id == "GO:0051607", ]
```

```
##                 gs_id            gs_description gs_pvalue
## GO:0051607 GO:0051607 defense response to virus   3.1e-13
##                                                                                                                                                                                                                                                                                   gs_genes
## GO:0051607 Apobec1,Bst2,Ccl5,Cd40,Cxcl10,Cxcl9,Ddx60,Dhx58,Eif2ak2,Fgl2,Gbp4,Ifih1,Ifit1,Ifit2,Ifit3,Ifitm1,Ifitm2,Ifitm3,Ifnb1,Il12b,Il12rb1,Il1b,Irf7,Isg15,Isg20,Itgax,Oas1a,Oas1c,Oas1g,Oas2,Oas3,Oasl1,Oasl2,Parp9,Pml,Rnasel,Rsad2,Rtp4,Slfn8,Slfn9,Stat1,Stat2,Trim12a,Tspan32,Zbp1
##            gs_de_count gs_bg_count Expected DE_count  z_score aggr_score
## GO:0051607          45         198    11.94       45 6.410062   2.939613
```

```
gs_heatmap(
  se = vst_alma,
  gtl = gtl_alma,
  geneset_id = "GO:0051607",
  cluster_columns = TRUE,
  anno_col_info = "condition"
)
```

##### 4.2.4 Comparing enrichment results

Imagine we were to compare the enrichment results from our study to another one where the same set of pathways is reported - or alternatively, on the same dataset but focusing on a different contrast.
We can replicate such a situation by shuffling the results in `res_enrich_a20`, and displaying the comparison with `gs_radar()`, `gs_summary_overview_pair()`, or `gs_horizon()`

```
# with more than one set
res_enrich_shuffled <- res_enrich_a20[1:60, ]
set.seed(42)
shuffled_ones <- sample(seq_len(60)) # to generate permuted p-values
res_enrich_shuffled$gs_pvalue <- res_enrich_shuffled$gs_pvalue[shuffled_ones]
# ideally, I would also permute the z scores and aggregated scores
gs_radar(res_enrich = res_enrich_a20,
         res_enrich2 = res_enrich_shuffled)
```

```
gs_summary_overview_pair(res_enrich = res_enrich_a20,
                         res_enrich2 = res_enrich_shuffled,
                         n_gs = 60)
```

```
gs_horizon(res_enrich = res_enrich_a20,
           compared_res_enrich_list = list(
             a20_shuffled = res_enrich_shuffled),
           n_gs = 40
          )
```

##### 4.2.5 Integrated reporting

The reporting functionality of `happy_hour()` can be called anytime also from the command line, focusing on a subset of genes and genesets of interest, which will be presented with additional detail in the output document.

```
happy_hour(
  gtl = gtl_alma,
  project_id = "a20-microglia",
  mygenesets = c(
    res_enrich_a20$gs_id[c(1:5, 11, 31)],
    # additionally selecting some by id
    "GO:0051963" # regulation of synapse assembly
  ),
  mygenes = c(
    "ENSMUSG00000035042",
    "ENSMUSG00000074896",
    "ENSMUSG00000035929"
  ),
  open_after_creating = TRUE
)
```

#### 4.3 Alternate entry point - using `edgeR` for the DE analysis

It is also possible to use `edgeR` to perform the steps before using `GeneTonic`.  
The chunk below reports an example of how to do so - imagine you start with a `DGE` object called `dge_alma`:

```
library("DEFormats")
library("edgeR")
dge_alma <- as.DGEList(readRDS("usecase_dds_alma.rds"))
```

We add the gene symbols for ease of readability…

```
dge_alma$genes$SYMBOL <- anno_df$gene_name[match(rownames(dge_alma$genes), anno_df$gene_id)]
```

… and perform a QL-based analysis, obtaining the `qlf` object at the end of it.

```
keep <- filterByExpr(dge_alma)
table(keep)
```

```
## keep
## FALSE  TRUE 
## 29148 12240
```

```
y <- dge_alma[keep, , keep.lib.sizes = FALSE]
y <- calcNormFactors(y)
conds <- dge_alma$samples$group
design <- model.matrix(~conds)
y <- estimateDisp(y, design)
# To perform quasi-likelihood F-tests:
fit <- glmQLFit(y, design)
qlf <- glmQLFTest(fit, coef = "condsko_del")
plotMD(qlf)
```

We would need to convert these objects back to the `DESeq2`-based objects.

```
convert_to_DESeqResults <- function(dgelrt, dge, FDR = 0.05) {
  tbledger <- as.data.frame(topTags(dgelrt, n = Inf))
  colnames(tbledger)[colnames(tbledger) == "PValue"] <- "pvalue"
  colnames(tbledger)[colnames(tbledger) == "logFC"] <- "log2FoldChange"
  colnames(tbledger)[colnames(tbledger) == "logCPM"] <- "baseMean"
  colnames(tbledger)[colnames(tbledger) == "SYMBOL"] <- "SYMBOL"
  
  # get from the logCPM to something on the scale of the baseMean
  tbledger$baseMean <- (2^tbledger$baseMean) * mean(dge$samples$lib.size) / 1e6
  
  # use the constructor for DESeqResults
  edger_resu <- DESeqResults(DataFrame(tbledger))
  edger_resu <- DESeq2:::pvalueAdjustment(edger_resu,
    independentFiltering = FALSE,
    alpha = FDR, pAdjustMethod = "BH"
  )
}

res_de_from_edgeR <- convert_to_DESeqResults(qlf, dge_alma)

summary(decideTestsDGE(qlf))
```

```
##        condsko_del
## Down            62
## NotSig       11923
## Up             255
```

```
summary(res_de_from_edgeR, alpha = 0.05)
```

```
## 
## out of 12240 with nonzero total read count
## adjusted p-value < 0.05
## LFC > 0 (up)       : 255, 2.1%
## LFC < 0 (down)     : 62, 0.51%
## outliers [1]       : 0, 0%
## low counts [2]     : 0, 0%
## (mean count < 0)
## [1] see 'cooksCutoff' argument of ?results
## [2] see 'independentFiltering' argument of ?results
```

```
dds_alma_from_edgeR <- as.DESeqDataSet(dge_alma)
```

Imagine we have already our `res_enrich` object, obtained in any of the ways specified above - then we can simply launch `GeneTonic` with these commands (utilizing the `anno_df` object from the examples above):

```
gtl_from_edgeR <- GeneTonic_list(
  dds = dds_alma_from_edgeR,
  res_de = res_de_from_edgeR,
  res_enrich = res_enrich_clusterprofiler,
  annotation_obj = anno_df
)
GeneTonic(gtl = gtl_from_edgeR)
```

### Session information

```
sessionInfo()
```

```
## R version 4.1.0 RC (2021-05-17 r80314)
## Platform: x86_64-apple-darwin17.0 (64-bit)
## Running under: macOS Mojave 10.14.6
## 
## Matrix products: default
## BLAS:   /System/Library/Frameworks/Accelerate.framework/Versions/A/Frameworks/vecLib.framework/Versions/A/libBLAS.dylib
## LAPACK: /Library/Frameworks/R.framework/Versions/4.1/Resources/lib/libRlapack.dylib
## 
## locale:
## [1] en_US.UTF-8/en_US.UTF-8/en_US.UTF-8/C/en_US.UTF-8/en_US.UTF-8
## 
## attached base packages:
## [1] parallel  stats4    stats     graphics  grDevices utils     datasets 
## [8] methods   base     
## 
## other attached packages:
##  [1] edgeR_3.33.7                limma_3.47.15              
##  [3] DEFormats_1.19.0            igraph_1.2.6               
##  [5] org.Mm.eg.db_3.13.0         magrittr_2.0.1             
##  [7] visNetwork_2.0.9            enrichR_3.0                
##  [9] msigdbr_7.4.1               fgsea_1.17.0               
## [11] tibble_3.1.2                gprofiler2_0.2.0           
## [13] clusterProfiler_3.99.2      dplyr_1.0.6                
## [15] apeglm_1.13.4               GeneTonic_1.3.8            
## [17] DT_0.18                     ideal_1.15.2               
## [19] pcaExplorer_2.17.1          bigmemory_4.5.36           
## [21] org.Hs.eg.db_3.13.0         pheatmap_1.0.12            
## [23] topGO_2.43.0                SparseM_1.81               
## [25] GO.db_3.13.0                AnnotationDbi_1.53.2       
## [27] graph_1.69.0                DESeq2_1.31.21             
## [29] SummarizedExperiment_1.21.3 Biobase_2.51.0             
## [31] MatrixGenerics_1.3.1        matrixStats_0.58.0         
## [33] GenomicRanges_1.43.4        GenomeInfoDb_1.27.13       
## [35] IRanges_2.25.11             S4Vectors_0.29.19          
## [37] BiocGenerics_0.37.6         knitr_1.33                 
## [39] job_0.2                     rmarkdown_2.8              
## 
## loaded via a namespace (and not attached):
##   [1] ps_1.6.0                 Rsamtools_2.7.2          foreach_1.5.1           
##   [4] rprojroot_2.0.2          crayon_1.4.1             MASS_7.3-54             
##   [7] backports_1.2.1          nlme_3.1-152             colourpicker_1.1.0      
##  [10] GOSemSim_2.17.2          rlang_0.4.11             XVector_0.31.1          
##  [13] callr_3.7.0              filelock_1.0.2           GOstats_2.57.0          
##  [16] BiocParallel_1.25.5      rjson_0.2.20             bit64_4.0.5             
##  [19] glue_1.4.2               rngtools_1.5             processx_3.5.2          
##  [22] UpSetR_1.4.0             shinyAce_0.4.1           shinydashboard_0.7.1    
##  [25] DOSE_3.17.2              tidyselect_1.1.1         XML_3.99-0.6            
##  [28] tidyr_1.1.3              GenomicAlignments_1.27.2 xtable_1.8-4            
##  [31] evaluate_0.14            ggplot2_3.3.3            cli_2.5.0               
##  [34] zlibbioc_1.37.0          rstudioapi_0.13          miniUI_0.1.1.1          
##  [37] bslib_0.2.5              fastmatch_1.1-0          BiocStyle_2.19.2        
##  [40] treeio_1.15.7            shiny_1.6.0              xfun_0.23               
##  [43] clue_0.3-59              pkgbuild_1.2.0           cluster_2.1.2           
##  [46] caTools_1.18.2           tidygraph_1.2.0          TSP_1.1-10              
##  [49] KEGGREST_1.31.2          expm_0.999-6             ggrepel_0.9.1           
##  [52] threejs_0.3.3            ape_5.5                  dendextend_1.15.1       
##  [55] shinyWidgets_0.6.0       Biostrings_2.59.4        png_0.1-7               
##  [58] withr_2.4.2              shinyBS_0.61             bitops_1.0-7            
##  [61] slam_0.1-48              ggforce_0.3.3            RBGL_1.67.0             
##  [64] plyr_1.8.6               GSEABase_1.53.2          coda_0.19-4             
##  [67] pillar_1.6.1             gplots_3.1.1             GlobalOptions_0.1.2     
##  [70] cachem_1.0.5             GenomicFeatures_1.43.8   GetoptLong_1.0.5        
##  [73] vctrs_0.3.8              ellipsis_0.3.2           generics_0.1.0          
##  [76] NMF_0.23.0               tools_4.1.0              munsell_0.5.0           
##  [79] tweenr_1.0.2             DelayedArray_0.17.13     fastmap_1.1.0           
##  [82] compiler_4.1.0           httpuv_1.6.1             rtracklayer_1.51.5      
##  [85] geneLenDataBase_1.27.0   pkgmaker_0.32.2          plotly_4.9.3            
##  [88] GenomeInfoDbData_1.2.6   gridExtra_2.3            lattice_0.20-44         
##  [91] AnnotationForge_1.33.6   utf8_1.2.1               later_1.2.0             
##  [94] BiocFileCache_1.99.8     jsonlite_1.7.2           scales_1.1.1            
##  [97] tidytree_0.3.3           genefilter_1.73.1        lazyeval_0.2.2          
## [100] promises_1.2.0.1         doParallel_1.0.16        checkmate_2.0.0         
## [103] goseq_1.43.0             cowplot_1.1.1            webshot_0.5.2           
## [106] downloader_0.4           survival_3.2-11          numDeriv_2016.8-1.1     
## [109] rsconnect_0.8.17         yaml_2.2.1               rintrojs_0.3.0          
## [112] htmltools_0.5.1.1        memoise_2.0.0            spelling_2.2            
## [115] BiocIO_1.1.2             locfit_1.5-9.4           seriation_1.2-9         
## [118] graphlayouts_0.7.1       viridisLite_0.4.0        digest_0.6.27           
## [121] assertthat_0.2.1         mime_0.10                commonmark_1.7          
## [124] rappdirs_0.3.3           emdbook_1.3.12           registry_0.5-1          
## [127] BiasedUrn_1.07           bigmemory.sri_0.1.3      RSQLite_2.2.7           
## [130] remotes_2.3.0            data.table_1.14.0        blob_1.2.1              
## [133] lpsymphony_1.19.0        labeling_0.4.2           splines_4.1.0           
## [136] Cairo_1.5-12.2           RCurl_1.98-1.3           hms_1.1.0               
## [139] colorspace_2.0-1         base64enc_0.1-3          BiocManager_1.30.15     
## [142] shape_1.4.5              aplot_0.0.6              sass_0.4.0              
## [145] Rcpp_1.0.6               bookdown_0.22            mvtnorm_1.1-1           
## [148] circlize_0.4.12          enrichplot_1.11.3        backbone_1.4.0          
## [151] fansi_0.4.2              IHW_1.19.0               R6_2.5.0                
## [154] grid_4.1.0               lifecycle_1.0.0          curl_4.3.1              
## [157] jquerylib_0.1.4          DO.db_2.9                Matrix_1.3-3            
## [160] qvalue_2.23.0            RColorBrewer_1.1-2       iterators_1.0.13        
## [163] stringr_1.4.0            htmlwidgets_1.5.3        hunspell_3.0.1          
## [166] polyclip_1.10-0          biomaRt_2.47.9           purrr_0.3.4             
## [169] crosstalk_1.1.1          shadowtext_0.0.8         ComplexHeatmap_2.7.11   
## [172] mgcv_1.8-35              patchwork_1.1.1          bdsmatrix_1.3-4         
## [175] codetools_0.2-18         bs4Dash_0.5.0            gtools_3.8.2            
## [178] prettyunits_1.1.1        dbplyr_2.1.1             gridBase_0.4-7          
## [181] gtable_0.3.0             DBI_1.1.1                dynamicTreeCut_1.63-1   
## [184] highr_0.9                httr_1.4.2               KernSmooth_2.23-20      
## [187] stringi_1.6.2            progress_1.2.2           reshape2_1.4.4          
## [190] farver_2.1.0             heatmaply_1.2.1          annotate_1.69.2         
## [193] viridis_0.6.1            rentrez_1.2.3            fdrtool_1.2.16          
## [196] Rgraphviz_2.35.0         magick_2.7.2             ggtree_2.99.0           
## [199] xml2_1.3.2               rvcheck_0.1.8            bbmle_1.0.23.1          
## [202] restfulr_0.0.13          geneplotter_1.69.0       Category_2.57.2         
## [205] bit_4.0.4                shinycssloaders_1.0.0    scatterpie_0.1.6        
## [208] ggraph_2.0.5             babelgene_21.4           pkgconfig_2.0.3
```
